# Paclitaxel- and vincristine-induced neurotoxicity and drug transport in sensory neurons

**DOI:** 10.1101/2023.02.07.527432

**Authors:** Christina Mortensen, Katherina C. Chua, Helen S. Hammer, Flemming Nielsen, Oliver Pötz, Åsa Fex Svenningsen, Deanna L. Kroetz, Tore Bjerregaard Stage

**Affiliations:** Clinical Pharmacology, Pharmacy, and Environmental Medicine, Department of Public Health, University of Southern Denmark, Odense, Denmark; Department of Bioengineering and Therapeutic Sciences, University of California, San Francisco, California, USA; Signatope GmbH, Reutlingen, Germany; Neurobiology Research Unit, Department of Molecular Medicine, University of Southern Denmark, Odense, Denmark; Department of Clinical Pharmacology, Odense University Hospital, Odense, Denmark

**Author notes:** Corresponding author for this work, Tore Bjerregaard Stage, J.B. Winsløws Vej 19, 2^nd^ floor, 5000 Odense C, Denmark, Twitter: @tstage1.

**Keywords:** Chemotherapy-induced peripheral neuropathy, paclitaxel, vincristine, induced pluripotent stem cells, sensory neurons, dorsal root ganglia, transient receptor potential vanilloid 1, efflux transporters, neurotoxicity, neuroprotection

## Abstract

Chemotherapy-induced peripheral neuropathy (CIPN) constitutes a significant health problem due to the increasing prevalence and the lack of therapies for treatment and prevention. Patients with CIPN primarily present with sensory symptoms, such as sensory disturbances that may progress to neuropathic pain in hands and feet. While pivotal for routine cancer treatment, paclitaxel and vincristine frequently cause CIPN and impact the quality of life among cancer patients and survivors. We utilized a model of human sensory neurons derived from induced pluripotent stem cells (iPSC-SNs) to provide mechanistic understanding of CIPN caused by paclitaxel and vincristine. The morphological phenotype of iPSC-SNs following paclitaxel exposure was characterized by retraction and thickening of axons while vincristine caused fragmentation and abolishment of axons. Both agents increased the mRNA expression of the pain receptor, transient receptor potential vanilloid (*TRPV1*), and highly induced neuronal damage, as measured by activating transcription factor 3 (*ATF3*) mRNA. iPSC-SNs express the efflux transporters, P-glycoprotein (P-gp, encoded by *ABCB1*) and multidrug resistance-associated protein 1 (MPR1, encoded by *ABCC1*). Inhibition of P-gp and MRP1 in iPSC-SNs exacerbated neurotoxicity of paclitaxel and vincristine respectively. We further show that pre-treatment with the P-gp inducer rifampicin alleviated chemotherapy-induced structural and transcriptional alterations in iPSC-SNs. iPSC-SNs are a valuable and robust model to study the role of efflux transporters and other mechanistic targets in CIPN. Efflux transporters play a critical role in CIPN pathogenesis as they regulate the disposition of chemotherapy to the peripheral nervous system.

## Introduction

Advances in cancer diagnosis and treatment have led to declining mortality rates over the last decades (1). Although contributing to increased cancer survival, treatment with chemotherapy is accompanied by multiple acute or long-term adverse effects. Chemotherapy-induced peripheral neuropathy (CIPN) is experienced by 2 out of 3 cancer patients undergoing neurotoxic chemotherapy (2). Neurotoxic chemotherapy includes taxanes (e.g., paclitaxel and docetaxel), vinca alkaloids (e.g., vincristine), platinum-based agents (e.g., oxaliplatin and cisplatin), proteasome inhibitors (e.g., bortezomib), and immunomodulatory agents (e.g., thalidomide) (3). CIPN presents with a broad range clinical symptoms depending on whether sensory, motor, or autonomic neurons are damaged by the specific type of neurotoxic chemotherapy (4). However, CIPN is primarily associated with sensory symptoms characterized by neuropathic pain and sensory disturbances in the extremities. CIPN can be partially reversible with recovery within several months to years after concluding treatment (5,6). In some cases, neurotoxic chemotherapy can cause irreversible nerve damage and CIPN symptoms that impact long-term quality of life among cancer survivors (7). According to clinical practice guidelines, there are currently no recommended agents for prevention of CIPN. Duloxetine is the only agent with moderate evidence for treatment of CIPN, yet with limited benefit for patients (8). Lack of effective treatment options for patients with intolerable CIPN symptoms leaves dose reductions, delays, discontinuation, or substitution as the only countermeasures. CIPN remains a major unsolved challenge within oncology due to lack of objective and easily applicable tools to accurately assess CIPN severity and to identify patients at increased risk of CIPN.

The pathogenesis of CIPN remains largely unknown. Multiple molecular mechanisms have been proposed; including, but not limited to, impaired axonal transport (9,10), mitochondrial dysfunction (11), neuroinflammation (12), altered neuronal excitability and expression of ion channels (13). Chemotherapeutic agents associated with CIPN have different targets of neurotoxicity, including the neuronal cell bodies of the dorsal root ganglion (DRG), the axons, or axonal components (e.g., microtubules, myelin sheaths, mitochondria, and ion channels) (14). Paclitaxel and vincristine, two antitubulin agents with opposite effects on microtubules, have both been proposed to exhibit neurotoxicity by targeting the distal nerve terminals on sensory neurons (15,16). Oxaliplatin on the other hand affects specific ion channels in cold-sensing sensory neurons, leading to neuronal hyperexcitability (17), while bortezomib primarily targets the myelin sheaths causing structural and functional alterations in sensory neurons (14). This explains why patients present with differences in symptom onset and clinical presentation when treated with distinct chemotherapies. However, heterogeneity in clinical presentation might also be observed within patient populations receiving the same agent (18).

The peripheral nervous system (PNS) is susceptible to circulating substances, such as chemotherapy, because it is not protected by the blood-brain barrier. Efflux transporters are critical for limiting the accumulation of toxic substances in the central nervous system (CNS) (19). Although their distribution has also been verified in the PNS (20), their role in drug disposition is unknown. Paclitaxel and vincristine are substrates for the major efflux transporter, P-glycoprotein (P-gp, encoded *ABCB1*). Vincristine is also a substrate for multidrug resistance-associated protein 1 (MRP1, encoded *ABCC1*) that mediate efflux of a broad range of drugs. Many commonly used drugs are P-gp inhibitors and thus, P-gp can be involved in drug-drug interactions. Chemotherapy has a narrow therapeutic index, and drug-drug interactions with chemotherapy may lead to severe toxicity (20–22).

Decades of research utilizing animal models have failed to translate into clinically effective therapeutic interventions. This may indicate that human model systems are warranted as they reflect human biology more closely. Human induced pluripotent stem cells (iPSCs) have become a powerful tool for providing an unlimited supply of human tissues to investigate CIPN pathophysiology and to screen for novel therapeutic targets. The purpose of this study was to study CIPN phenotypes using human sensory neurons derived from iPSCs (iPSC-SNs). Specifically, we investigated morphological and transcriptional alterations upon exposure to the neurotoxic chemotherapeutic agents, paclitaxel, and vincristine. Furthermore, we assessed the impact of inhibition or induction of efflux transporters on chemotherapy-induced neurotoxicity in iPSC-SNs.

## Methods

### Differentiation of sensory neurons

Human iPSCs from a healthy female donor (A18945, ThermoFisher, Roskilde, Denmark, https://hpscreg.eu/cell-line/TMOi001-A) were differentiated into sensory neurons as previously described (23). iPSCs were maintained in mTeSR1 medium (85850, StemCell Technologies, Vancouver, BC, Canada) on Matrigel (diluted according to manufacturer’s instructions, 354277, Corning, NY, USA) at a minimum density of 50,000 cells/cm^2^. The medium was changed daily, and iPSCs were clump-passaged at 70-80% confluency using Accutase (00455556, ThermoFisher). At the final passage before differentiation, iPSCs were seeded as single cells onto Matrigel-coated 6-well plates (354277, Corning, NY, USA) using Accutase. iPSCs were maintained in mTeSR1 medium (85850, StemCell Technologies, Vancouver, BC, Canada) until >90% confluency. After 72 hours, the differentiation into sensory neurons was initiated using KSR medium which contained 82% KnockOut DMEM (10829018, ThermoFisher), 15% KnockOut Serum Replacement (10828028, ThermoFisher), 1% GlutaMAX (35050038, ThermoFisher, Paisley, Scotland, UK), 1% MEM non-essential amino acids (11140035, ThermoFisher, Paisley, Scotland, UK), 1% Penicillin-Streptomycin (P4333, Sigma-Aldrich, Søborg, Denmark) and 0.1 mM β-mercaptoethanol (31350010, ThermoFisher, Paisley, Scotland, UK). The two small molecule inhibitors LDN193189 (0.1 μM, S7507, Selleck Chemicals) and SB431542 (10 μM, S1067, Selleck Chemicals) were added on days 0-5. Three additional small molecule inhibitors consisting of CHIR99021 (3 μM, S1263, Selleck Chemicals), SU5402 (10 μM, S7667, Selleck Chemicals) and DAPT (10 μM, S2215, Selleck Chemicals) were added on days 2-11. The medium was changed daily between days 0-12, and N2 medium was incremented by 25% every other day starting from day 4. N2 medium consisted of 50% DMEM/F-12 (11320033, ThermoFisher) and 50% Neurobasal (21103049, ThermoFisher) with 1% N2 Supplement (17502048, ThermoFisher), 1% B27 Supplement (17504001, ThermoFisher) and 1% Penicillin-Streptomycin. Immature sensory neurons were dissociated and seeded as single cells onto plates coated with poly-L-ornithine hydrobromide (20 µg/mL, P3655, Sigma-Aldrich), laminin (10 µg/mL, 23017015, ThermoFisher), and fibronectin (2 µg/mL, F1141, Sigma-Aldrich) at a density of 150,000 viable cells/cm^2^. Immature iPSC-SNs were maintained in N2 medium supplemented with human growth factors (25 ng/mL NGF-β, 450-01; BDNF, 450-02; GDNF, 450-10; NT-3, 450-03; Peprotech, Cranbury, NJ, USA), and 0.2 mM L-ascorbic acid (A4403, Sigma-Aldrich). To remove fibroblasts from the culture, cells were exposed to freshly prepared Mitomycin-C (1 µg/mL, M4287, Sigma-Aldrich) for 2 hours on day 14. The medium was completely replaced on day 16 and on days 35-45 where experiments was performed. On all other days, 50% of the medium was changed every 3-4 days. To improve cell attachment, laminin (1 µg/mL) was added to the medium once a week.

### Compound preparations

Paclitaxel (T7402,), 5-fluorouracil (5-FU, FF2627), valspodar (SML0572), MK-571 (M7571), and rifampicin (R3501, Sigma-Aldrich) were purchased from Sigma-Aldrich (Søborg, Denmark) and dissolved in dimethyl sulfoxide (DMSO). Vincristine (V8388, Sigma-Aldrich) was prepared on ice and in the dark with phosphate-buffered saline containing calcium and magnesium (PBS, Sigma, D8662). The final concentration of DMSO or PBS was 0.2% in each experiment. Mature iPSC-SNs were exposed to paclitaxel, vincristine, and 5-fluorouracil for 48 hours. Clinically relevant concentrations of chemotherapy were selected to ensure translation of results (3).

### Repeated exposure

To assess single versus repeated exposure, mature iPSC-SNs were exposed to one or two concentrations of paclitaxel in parallel. After a single exposure of 0.01, 0.1, and 1 µM paclitaxel for 48 hours; one plate was fixed and for the other plate, the medium was completely removed and replaced with fresh and paclitaxel-free N2 medium. On day 3, 50% of the medium was changed and on day 5, iPSC-SNs were exposed to a second concentration of paclitaxel for 48 hours.

### Inhibition and induction of efflux transporters

To allow binding to efflux transporters, iPSC-SNs were pre-exposed to 4 μM valspodar or 4 μM MK-571 for 1 hour before exposure to paclitaxel or vincristine for 48 hours.

To allow transcription of efflux transporters, iPSC-SNs were pre-exposed to 10 µM rifampicin for 48 hours. Following induction, all medium was removed and iPSC-SNs was exposed to paclitaxel or vincristine for 48 hours.

### Immunolabeling

Mature iPSC-SNs were fixed by diluting 16% paraformaldehyde (28906, ThermoFisher) 1:4 into the cell medium for 10 minutes. After two washing steps with PBS containing calcium and magnesium, iPSC-SNs were permeabilized with 0.25% Triton X-100 for 15 minutes. Unspecific binding was subsequently blocked using 1% bovine serum albumin (A9418, Sigma-Aldrich) for 1 hour. iPSC-SNs were labeled with peripherin (1:200, SC-377093, Santa Cruz) and TRPV1 (1:100, ACC-030, Alomone Labs) overnight at 4°C. The following day, iPSC-SNs were labeled with Alexa Fluor 488-conjugated anti-rabbit (1:400, A21448, ThermoFisher) and Rhodamine Red-X-conjugated anti-mouse (1:600, 715-295-150, Jackson ImmunoResearch) for 1 hour at room temperature. iPSC-SNs were subsequently washed twice with PBS and counterstained with 10 μM DAPI (D9542, Sigma-Aldrich) to visualize nuclei. To minimize cell detachment, 100% of medium was only removed at the final washing step after fixation and before addition of primary and secondary antibodies as well as DAPI. For all other steps, 10% of the washing or blocking buffer remained in each well. For immunolabeling performed in 24-well plates, iPSC-SNs were stored in PBS, and images were acquired using ImageXpress Pico (Molecular Devices, San Jose, CA, USA). For immunolabeling performed in 2-well Lab-Tek Chambers slides (177429, ThermoFisher), drops of Mowiol 4-88 (81381, Sigma-Aldrich) were added and cover glasses (613-0137, VWR) were placed on top. Images were acquired on the Leica DMI 4000B Inverted Microscope (Leica Microsystems, Wetzlar, Germany).

### Neurotoxicity assessment

We assessed neurotoxicity by measuring the area of the neuronal network and the number of axons emanating from each ganglion using Sholl analysis (ImageJ software 2.0.0). We obtained three images per triplicate conditions from 2-3 independent differentiations. All images were converted to 8-bit, and an appropriate threshold specific for each differentiation was applied. The center of the ganglion was defined using the straight-line tool, and Sholl analysis was performed for all ganglia in each image (**Figure S1**). The number of ganglia with >30 axons was subsequently counted for each chemotherapy concentration and is presented relative to the total number of ganglia counted for each chemotherapy concentration.

### MIPAR analysis

Quantitative imaging analysis was completed using a custom-built algorithm using MIPAR™ software as previously described (24,25). In short, three individual fields were imaged for each experimental well. Images were batch processed and the amount of peripherin-labeled neurite networks were quantified while minimizing non-specific labeling (i.e., neurite length). Quantifications from experimental wells were normalized accordingly to the control wells from the same differentiation.

### Fluorescence-activated cell sorting

Direct flow cytometric analysis of Ki-67 was performed on day 12 iPSC-SNs as previously described [26]. iPSC-SNs were analyzed along with SH-SY5Y neuroblastoma cells as positive control. Unlabeled controls were included for measuring the autofluorescence and for localizing the negative population.

iPSC-SNs and SH-SY5Y cells were washed once in PBS without calcium and magnesium (PBS÷), and subsequently dissociated with Accutase or TrypLE Express respectively (5 min, 37°C). Enzymatic activity was neutralized, and the cell suspensions were centrifuged (iPSC: 800 rpm; SH-SY5Y: 1000 rpm, 5 min, 4°C) and resuspended in PBS÷. For obtaining single cells, the cell suspensions were filtered through a 40 µm cell strainer (Corning, 352340). The total cell number were determined using Countess II FL Automated Cell Counter (Invitrogen) to ensure the presence of at least 200,000 viable cells per 100 µL cell suspension. The single cell suspensions were fixed with 1% PFA (15 min), permeabilized in 0.7% Tween-20 and blocked in 10% goat serum (30 min, 16210072, ThermoFisher, New Zealand). The single cell suspensions were incubated with primary antibodies diluted in dilution buffer containing 10% goat serum, 0.28% Tween and 1% BSA in PBS for at least 30 min at 4°Cin the dark. The primary conjugated antibodies include FITC-conjugated monoclonal Ki67 (1:100, eBioscience™, 11-5698-82) which was titrated prior to the experiments. After incubation with primary antibody, the cells were washed once in PBS÷ by centrifugation (300 g, 4°C, 5 min) and subsequently resuspended in PBS÷ with 1% BSA. For washing steps, the supernatant was aspired before adding any new solution, allowing a minor volume of liquid (approx. 100 µL to remain). However, the repeated washing leads to a noticeable amount of cell loss and therefore, an initial cell concentration of 2.0 · 10^6^ cells/mL is required to capture enough events within the population gate and ensure analytical quality.

Flow cytometric analysis was performed on LSR II flow cytometer (BD Biosciences, San Jose, CA, USA) immediately after labeling, and data were acquired with BD FACSDiva Software 8.0.1. The cells of interest were gated based on their forward scatter (FSC) and side scatter (SSC) by excluding dead cells and debris. To avoid excluding non-spherical cells of interest, doublet discrimination was not performed. Histogram profiles of fluorescence intensity were created for the FSC vs. SSC gated population, and fluorescence-detecting gates were established by acquiring data for the unlabeled control. The percentage of positive cells for the specific marker is determined by the shift in fluorescence intensity of the labeled cells compared to unlabeled cells.

### Quantitative real-time polymerase chain reaction

Total RNA was extracted from iPSC-SNs seeded in 12-well plates using RNeasy Mini Kit (74106, Qiagen AB, Stockholm, Sweden) with DNase digestion (79256, Qiagen). cDNA was synthesized using the High-Capacity cDNA Reverse Transcription Kit with RNase Inhibitor (4374966, Applied Biosystems, Foster City, CA, USA). Quantitative real-time polymerase chain reaction (qPCR) was performed by transferring 10 ng cDNA template to each well in a 96-well reaction plate that contained target-specific TaqMan probes (**Table S1**) and TaqMan Universal Master Mix II (final volume of 10 µL). A no reverse transcriptase control and a no template control were included for all target-specific probes. qPCR reactions were performed using StepOne Plus RT-PCR equipment (ThermoFisher, Carlsbad, CA, USA) according to the manufacturer’s instructions for 40 cycles. Data were analyzed using the comparative C_t_ method (26) and involved normalization to the housekeeping gene glyceraldehyde-3-phosphate (GAPDH).

### Quantification of intracellular paclitaxel concentrations by liquid chromatography and tandem mass spectrometry

Matured iPSC-SNs seeded in 12-well plates were exposed to 0.1, 1, and 10 µM paclitaxel for 1 hour. Cells were subsequently washed with PBS containing calcium and magnesium, and lysed using radioimmunoprecipitation assay buffer (89900, ThermoFisher) for 10 minutes on ice. Cell lysates were collected using a cell scraper, vortexed thoroughly for 1 minute, and centrifuged at 14,000 rpm for 10 min at 4°C. The supernatant was carefully collected and stored at −80°C until further analysis.

The paclitaxel concentration in the cell lysate was determined by use of LC-MS/MS as previously described [1]. Briefly, a volume of 100 μL cell lysate and 100 μL acetonitrile was transferred into a 1.5 mL polypropylene microtube (Sarsted, Nümbrecht, Germany), vortex mixed for 5 minutes and centrifuged at 15,000 *g* for 15 minutes. A volume of 10 μL of the supernatant was injected onto a LC-MS/MS system consisting of a Dionex Ultimate 3000 UHPLC system and a TSQ Vantage Triple Quadropole Mass Spectrometer with Heated Electrospray Ionization (Thermo Scientific, San Jose, CA). Calibration curves ranging from 25 nM to 3,000 nM as well as quality control samples were included in each batch of analysis. The within-batch imprecision of the method was < 8%. Limit of detection was 1 nM and lower limit of quantification was 5 nM. Finally, the paclitaxel concentrations were adjusted for protein concentration in each well as assessed using bicinchoninic acid (BCA) assay according to the manufacturer’s instructions (23227, ThermoFisher, Waltham, MA, USA),

### Quantification of drug transporters by immunoaffinity liquid chromatography and tandem mass spectrometry

The protein quantification was conducted as previously described (27). Briefly, prior to analysis, the total protein amount was determined via BCA assay, and 150 µg protein was proteolyzed using trypsin (Pierce Trypsin Protease, MS-grade; Thermo Scientific, Waltham, MA, USA). Surrogate peptides and internal standard peptides were precipitated using triple X proteomics (TXP) antibodies. Peptides were then eluted and quantified using a modified version of the previously described LC–MS methods (28,29): There, 6 min and 10 min parallel reaction monitoring (PRM) methods are described (UltiMate 3000 RSLCnano and PRM–QExactive Plus; Thermo Scientific, Waltham, MA, USA). Raw data were processed using TraceFinder 4.1 (Thermo Scientific, Waltham, MA, USA). Peptide amounts were calculated by forming the ratios of the integrated peaks of the endogenous peptides and the isotope-labeled standards. Quantities of proteins were reported normalized as fmol per µg extracted protein.

### Statistics

R (version 4.0.2; R Statistical Foundation for Statistical Computing, Vieanna, Austria) was used for data analysis and visualization. Neurotoxicity parameters were calculated relative to the mean of the respective vehicle control (DMSO or PBS) for each individual differentiation.

## Results

### Characterization of iPSC-derived sensory neurons

Sensory neurons were differentiated from a healthy iPSC donor as previously described (23). Briefly, iPSCs were differentiated into immature sensory neurons using a combination of small molecule inhibitors for 12 days followed by maturation with growth factors and ascorbic acid for additional 23-33 days. The differentiation was initiated on day 0 when iPSCs reached >90% confluency (**Figure 1A**). The cells slowly self-organized into clusters (**Figure 1B**) which further developed into round and 3D-like structures with numerous neurites (**Figures 1C and 1D**). After re-seeding on day 12, we observed that the immature sensory neurons were pseudounipolar (**Figure 1E**), a unique characteristic for neurons of the DRG. The low expression (15.2%) of the proliferation marker Ki-67 indicates that the majority iPSC-derived cells were postmitotic on day 12 (**Figure S2**). The mature sensory neurons had their cell bodies arranged in ganglia-like structures and developed comprehensive neuronal networks (**Figure 1F**). A group of sensory neurons were localized individually outside the ganglia-like structures which may suggest presence of different sensory neuron subtypes (**Figure S3**). The mature sensory neurons expressed the PNS marker, peripherin (**Figure 2**). The transient receptor potential vanilloid 1 (TRPV1), which is a key mediator of pain, was ubiquitously expressed in ganglia-like structures (**Figure 2**). Collectively, the resulting iPSC-SNs display a characteristic DRG morphology and express canonical DRG markers.

**Figure 1.**
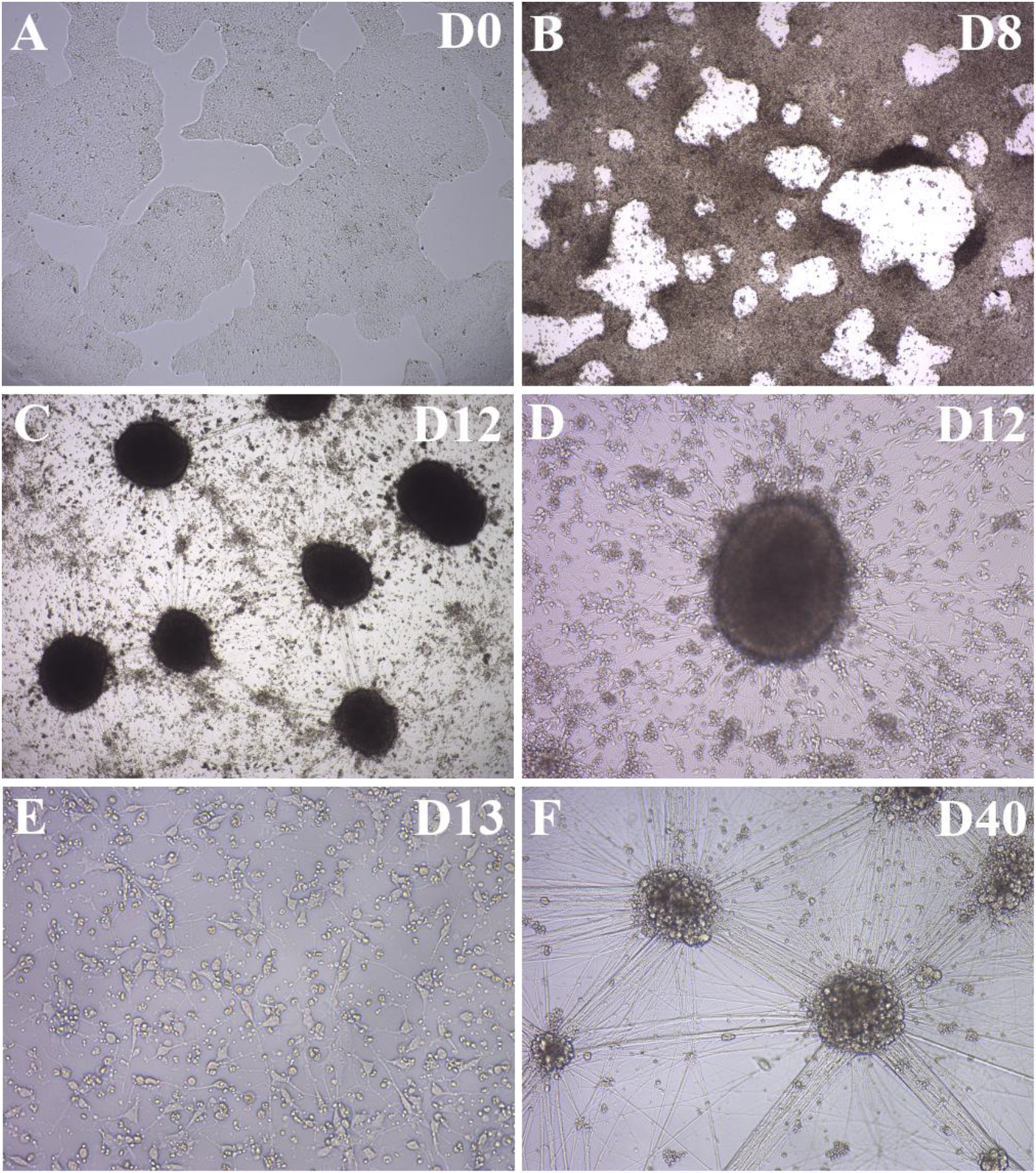
Overview of the differentiation of human iPSCs into sensory neurons. During the differentiation, the morphology changes significantly from iPSC colonies with well-defined borders to sensory neurons organized in ganglia-like structures with a large neuronal network. Images were acquired with VisiScope IT414. The 4X objective was used for images A-C, 10X for D and F, and 20X for E. Abbreviations: D, day; iPSC, induced pluripotent stem cell.

**Figure 2.**
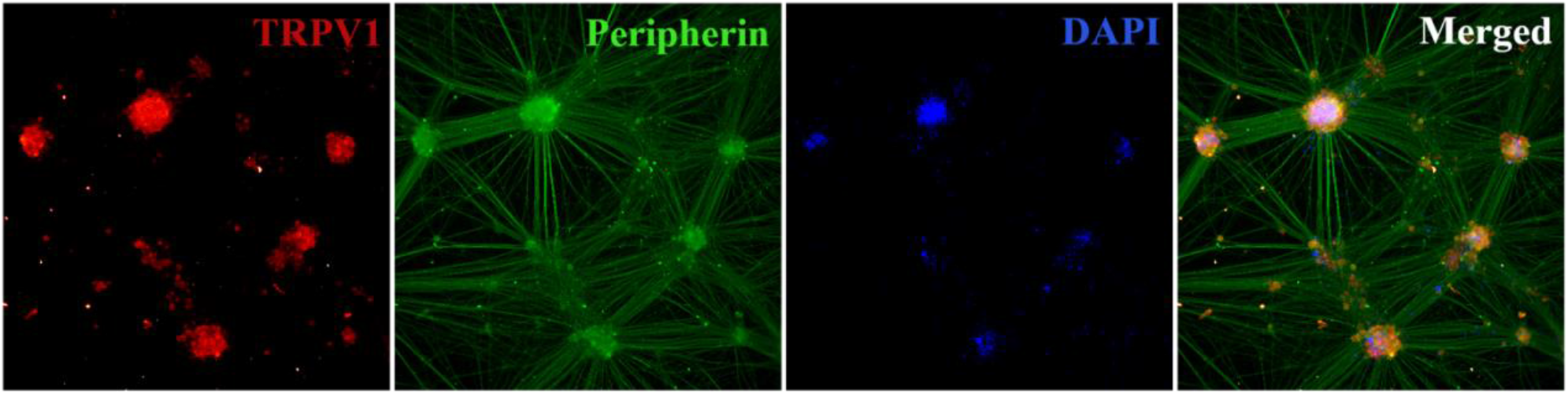
iPSC-derived sensory neurons (iPSC-SNs) express canonical dorsal root ganglia markers. Immunolabeling of day 39 iPSC-SNs was performed with TRPV1, peripherin, and DAPI. Images were obtained using ImageXpress Pico with the 10X objective. Abbreviations: DAPI, 4′,6-diamidino-2-phenylindole; TRPV1, transient receptor potential vanilloid 1.

### Paclitaxel causes retraction and thickening of axons and reduces the neuronal network

Exposure to paclitaxel caused axons of iPSC-SNs to retract (white arrows) and reduced the neuronal network in a concentration-dependent manner (**Figure 3**). Quantification of peripherin-labeled images showed that paclitaxel reduced the number of axons projecting from each ganglion (**Figure 5A**). The number of ganglia with >30 axons were reduced from 97% to 54% upon exposure to the highest concentration of 10 µM paclitaxel (**Figure 6A**). Repeated exposure of paclitaxel was observed to further exacerbate neurotoxicity in iPSC-SNs compared to single exposure (**Figures S4 and S5**). We assessed sensory neuronal accumulation of paclitaxel and found that it increased exponentially with increasing paclitaxel concentrations (**Figure 7**). This may suggest saturation of active efflux transport and indicates that active efflux transporters are involved in regulating the intracellular accumulation of paclitaxel into iPSC-SNs.

**Figure 3.**
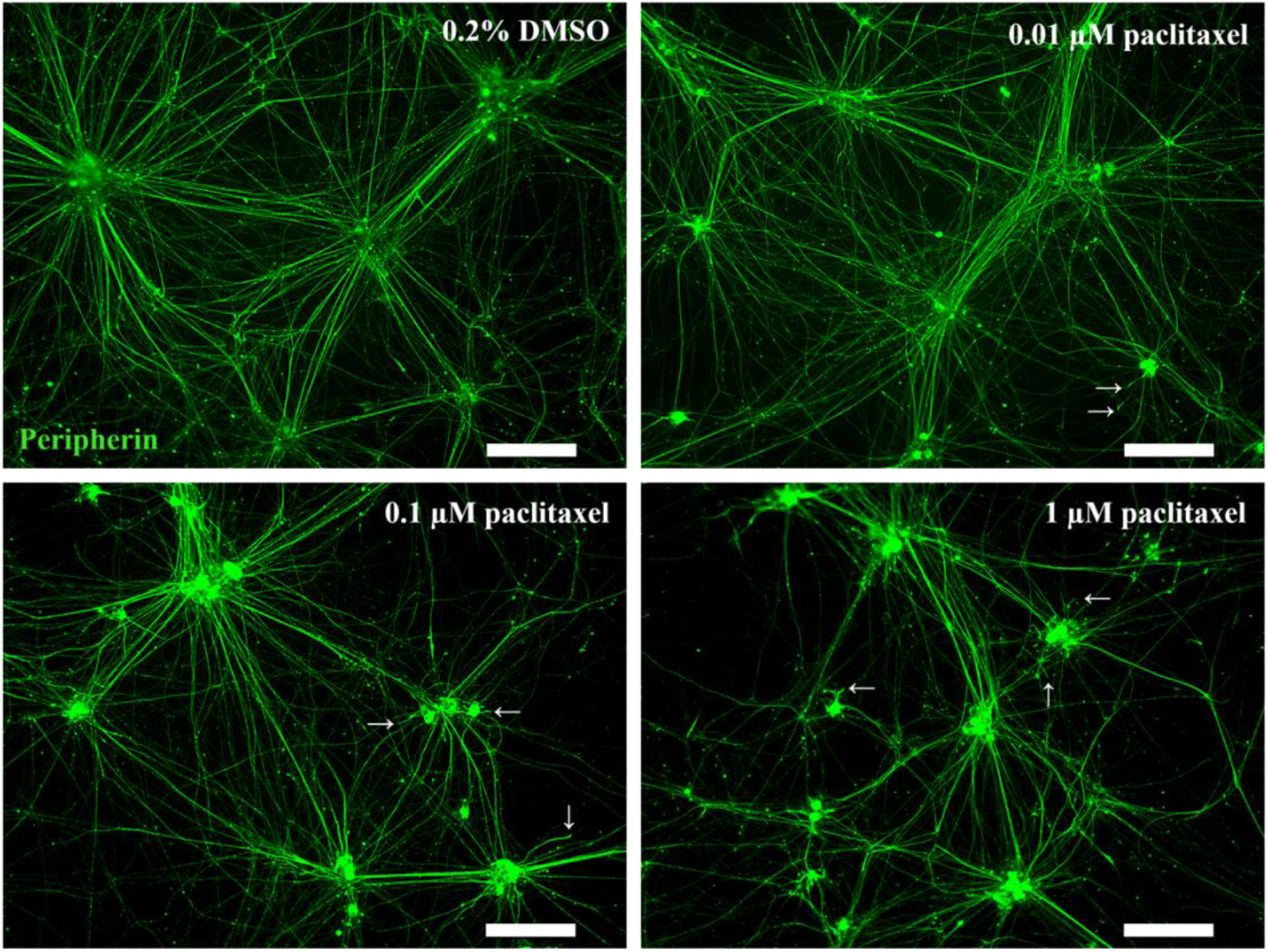
Paclitaxel causes axonal retraction (white arrowheads) and reduces the neuronal network in a concentration-dependent manner. Mature iPSC-derived sensory neurons were exposed to vehicle (0.2% DMSO) and indicated concentrations of paclitaxel for 48 hours followed by fixation and immunolabelingwith peripherin. Images were captured using the Leica DMI 4000B microscope with a 10X objective. Scale bars (white boxes) represent 200 μm. Abbreviations: DMSO, dimethyl sulfoxide; iPSC, induced pluripotent stem cell.

### Vincristine causes fragmentation and abolishment of the neuronal network

Exposure to vincristine caused fragmentation and abolishment of the neuronal network (**Figure 4**), even at the lowest concentration of 0.01 µM. Quantification of the labeled images demonstrated a substantial reduction of the number of axons and the area of the neuronal network upon increasing concentrations to vincristine (**Figure 5B**). We found that vincristine caused a concentration-dependent reduction of the number of ganglia with >30 axons (**Figure 6B**).

**Figure 4.**
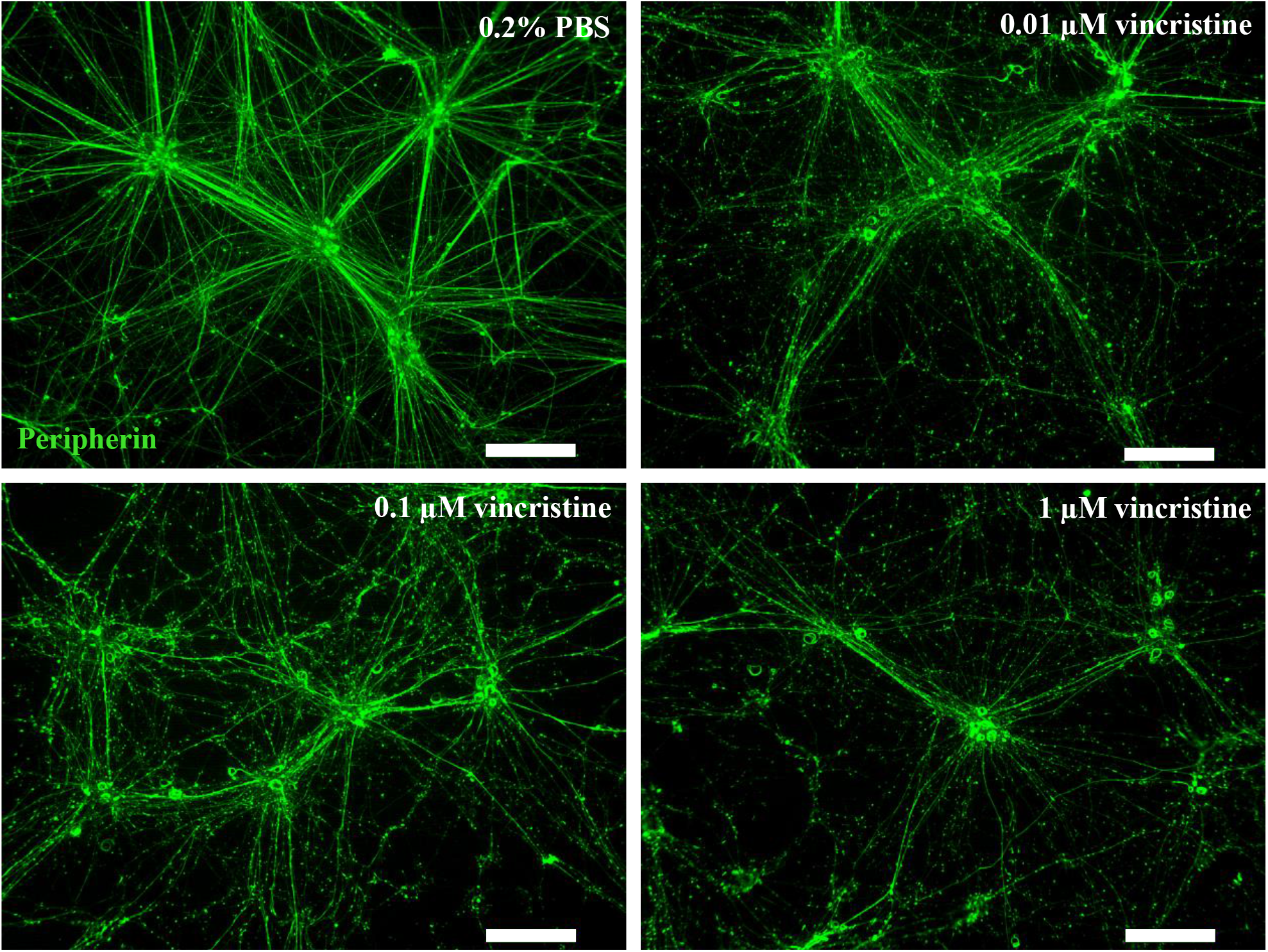
Vincristine causes axonal fragmentation of the neuronal network in a concentration-dependent manner. Mature iPSC-derived sensory neurons were exposed to vehicle (0.2% PBS) and indicated concentrations of vincristine for 48 hours followed by fixation and immunolabeling with peripherin. Images were captured using the Leica DMI 4000B microscope with a 10X objective. Scale bars represent 200 μm. Abbreviations: iPSC, induced pluripotent stem cell; PBS, phosphate-buffered saline.

**Figure 5.**
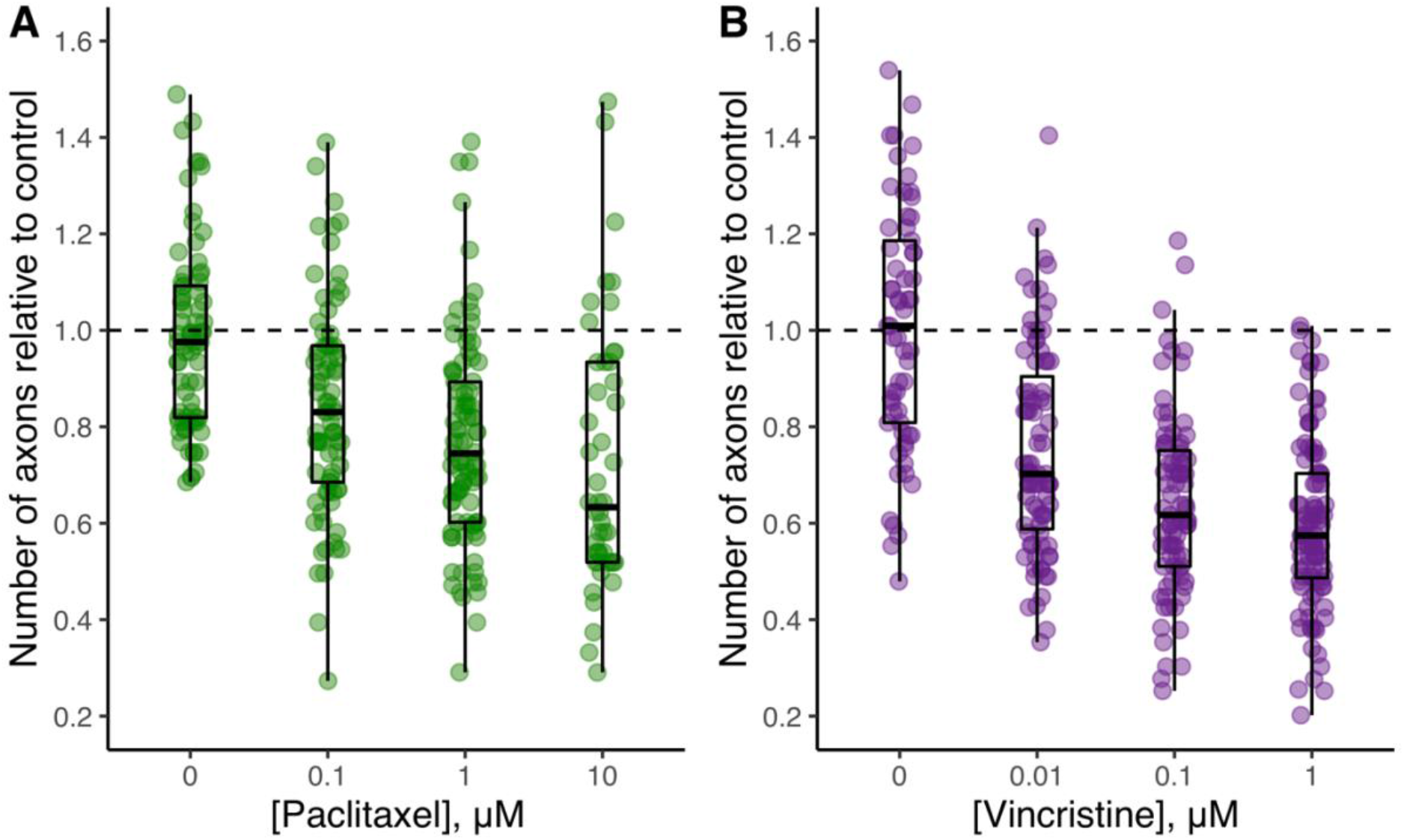
Paclitaxel and vincristine reduce the number of axons emanating from each ganglion. iPSC-derived sensory neurons were treated for 48 hours with vehicle (0.2% DMSO or 0.2% PBS) and the indicated concentrations of paclitaxel or vincristine. The results are shown relative to the vehicle from each differentiation (n=2 for paclitaxel, n=2 for vincristine).

**Figure 6.**
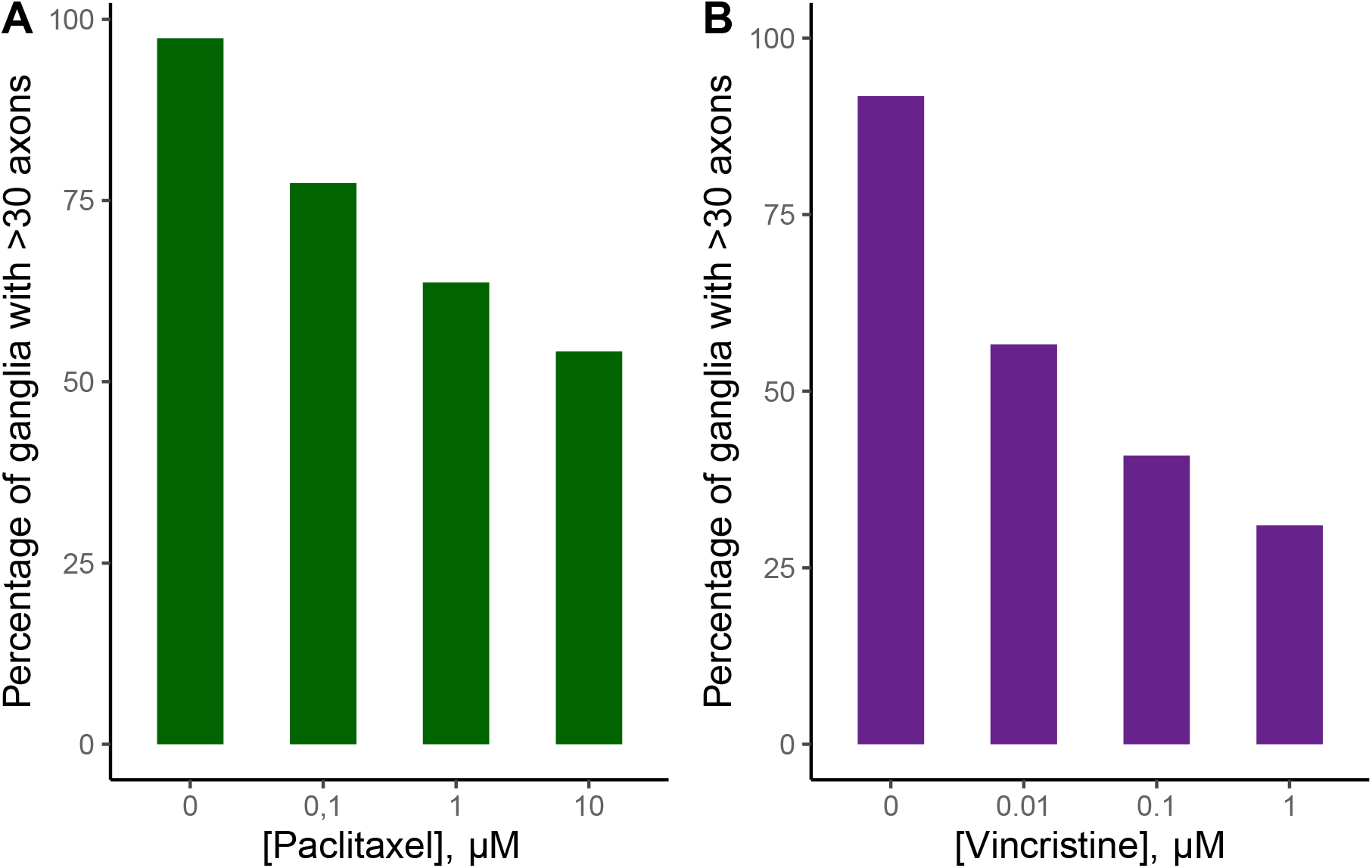
Paclitaxel and vincristine cause a dose-dependent reduction of the number of ganglia with >30 axons. The number of ganglia with >30 axons is presented relative to the total number of ganglia analyzed for each chemotherapy concentration (n=2 for paclitaxel, n=2 for vincristine).

**Figure 7.**
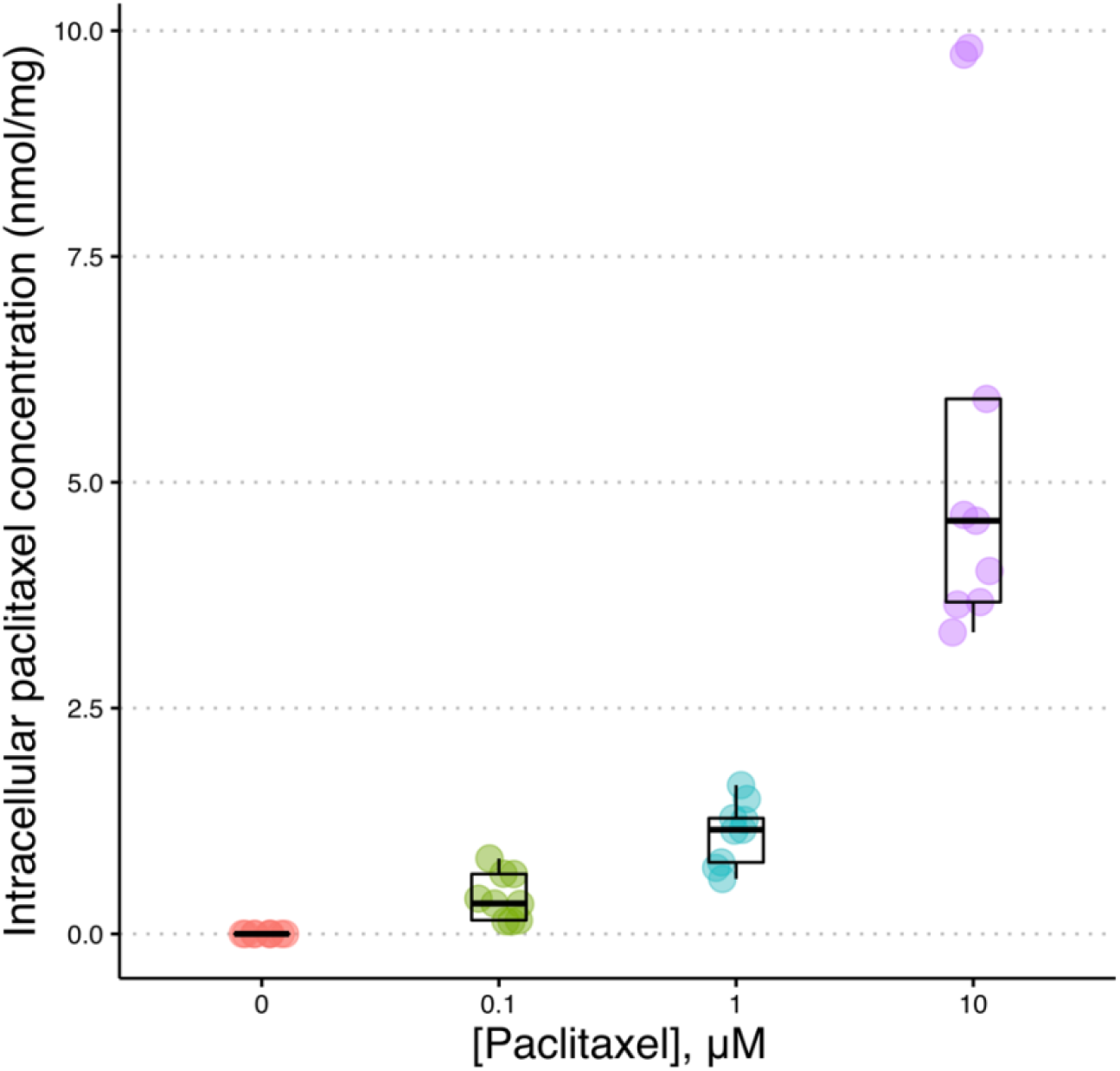
Paclitaxel accumulates in iPSC-derived sensory neurons with increasing concentrations. Mature iPSC-derived sensory neurons were exposed to vehicle (0.2% DMSO) or indicated concentrations of paclitaxel for 1 hour. Intracellular paclitaxel concentrations were measured in cell lysates using LC-MS/MS (n=3). Abbreviations: iPSC, induced pluripotent stem cell; LC-MS/MS, liquid chromatography and tandem mass spectrometry.

### Transcriptional alterations associated with paclitaxel and vincristine exposure

Mature iPSC-SNs were exposed to paclitaxel and vincristine to assess alterations of selected genes. Activating transcription factor 3 (*ATF3*) is considered a marker of damage in DRG and other tissues (30). We found that *ATF3* mRNA levels were highly upregulated in iPSC-SNs following exposure to both agents, though vincristine provoked the most dramatic upregulation (**Figure 8**). Exposure to paclitaxel and vincristine increased the expression of *TRPV1* in iPSC-SNs (**Figure 8**). Both agents also caused an induction of *ABCB1* that encodes the efflux transporter, P-glycoprotein (**Figure 8**). Vincristine, but not paclitaxel, caused an induction of *ABCC1* that encodes the efflux transporter MRP1 (**Figure 8**). The negative control (100 µM 5-FU) did not affect the neuronal network or increase *TRPV1* and *ATF3* mRNA levels. However, 5-FU downregulated *ABCB1* as well as *ABCC1* mRNA levels (**Figure S6**).

**Figure 8.**
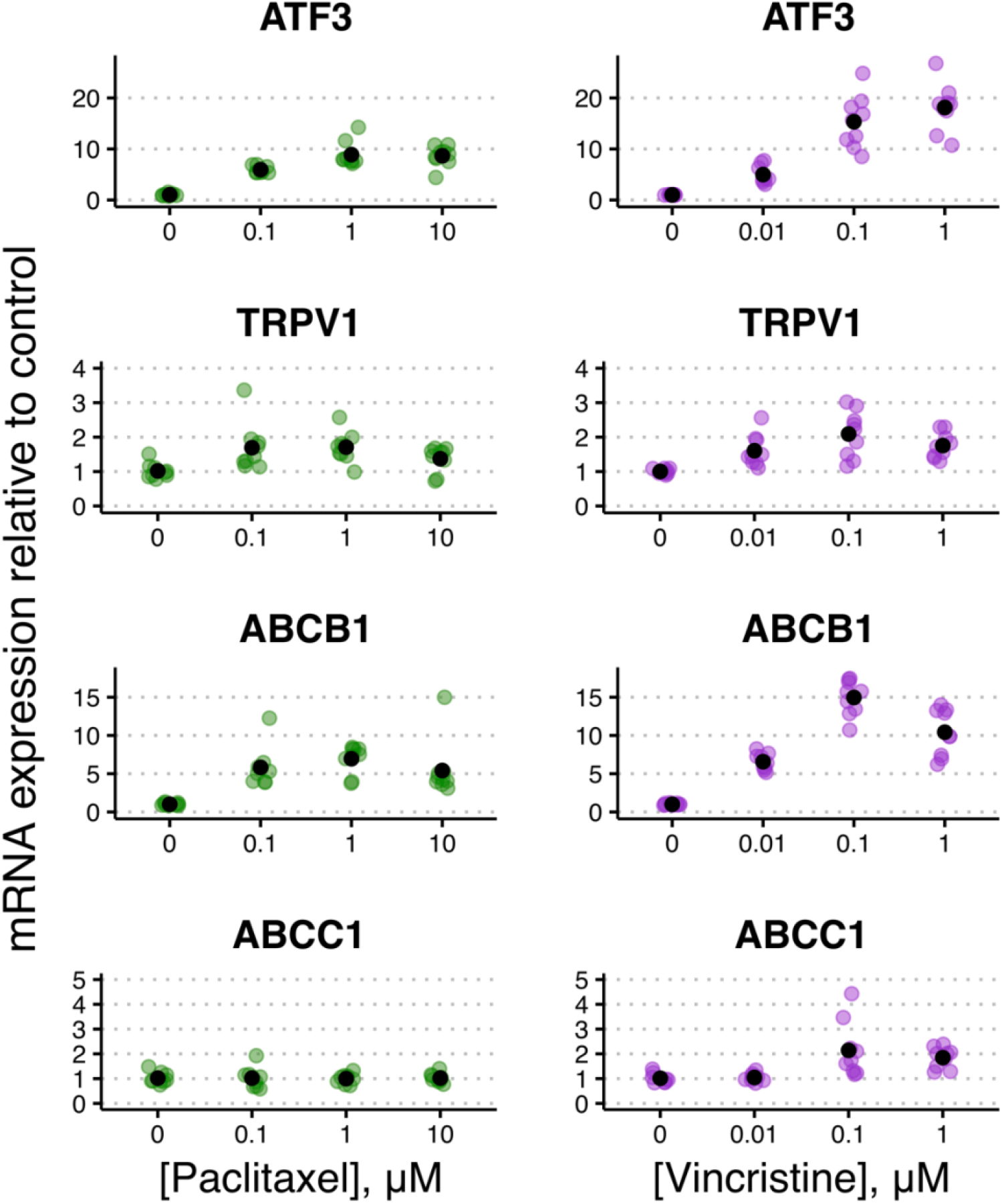
The effect of paclitaxel and vincristine on the mRNA expression of *ATF3, TRPV1, ABCB1*, and *ABCC1*. iPSC-derived sensory neurons were treated for 48 hours with vehicle (0.2% DMSO or 0.2% PBS) and the indicated concentrations of paclitaxel and vincristine. The mRNA level is presented relative to the vehicle from each differentiation. The black dot represents the mean value of 3 wells per condition from the 3 independent differentiations. Abbreviations: ABC, ATP-binding cassette; ATF3, activating transcription factor 3; DMSO, dimethyl sulfoxide; induced pluripotent stem cell-derived sensory neurons; PBS, phosphate-buffered saline; TRPV1, transient receptor potential vanilloid 1.

### Inhibition of P-gp and MRP1 might exacerbate paclitaxel- and vincristine neurotoxicity

The protein expression of relevant drug transporters was assessed using LC-MS/MS. We first verified that iPSC-SNs express the ABC transporters, P-gp (*ABCB1*), MRP1 (*ABCC1*), and breast cancer resistance protein (*ABCG2*, **Table S2**). To assess the role of P-gp on neurotoxicity, iPSC-SNs were pre-exposed to a P-gp inhibitor (valspodar) for 1 hour before exposure to paclitaxel. Our findings indicated that inhibition of P-gp exacerbates the neurotoxicity of paclitaxel, as observed by reduction of the neuronal network and an increased number of axonal retractions (**Figures 9 and S7**). Sholl analysis demonstrated that P-gp inhibition significantly reduced the number of axons for 0.01 μM and 0.1 μM paclitaxel compared to the DMSO control (**Figure 9A**). Interestingly, P-gp inhibition did not exacerbate neurotoxicity in iPSC-SNs at paclitaxel concentrations of 1 μM and 10 μM. Similar to axon number, neurite length was only reduced by P-gp inhibition at 0.01 µM paclitaxel (**Figure S8A**). Additionally, we assessed the role of MRP1 on neurotoxicity by pre-exposing iPSC-SNs to a MRP1 inhibitor (MK-571) prior to vincristine exposure. We showed that inhibition of MRP1 increased axonal fragmentation and substantially reduced the number of axons projecting from each ganglion as well as the neurite length (**Figures 9B, S8B, and S9**).

**Figure 9.**
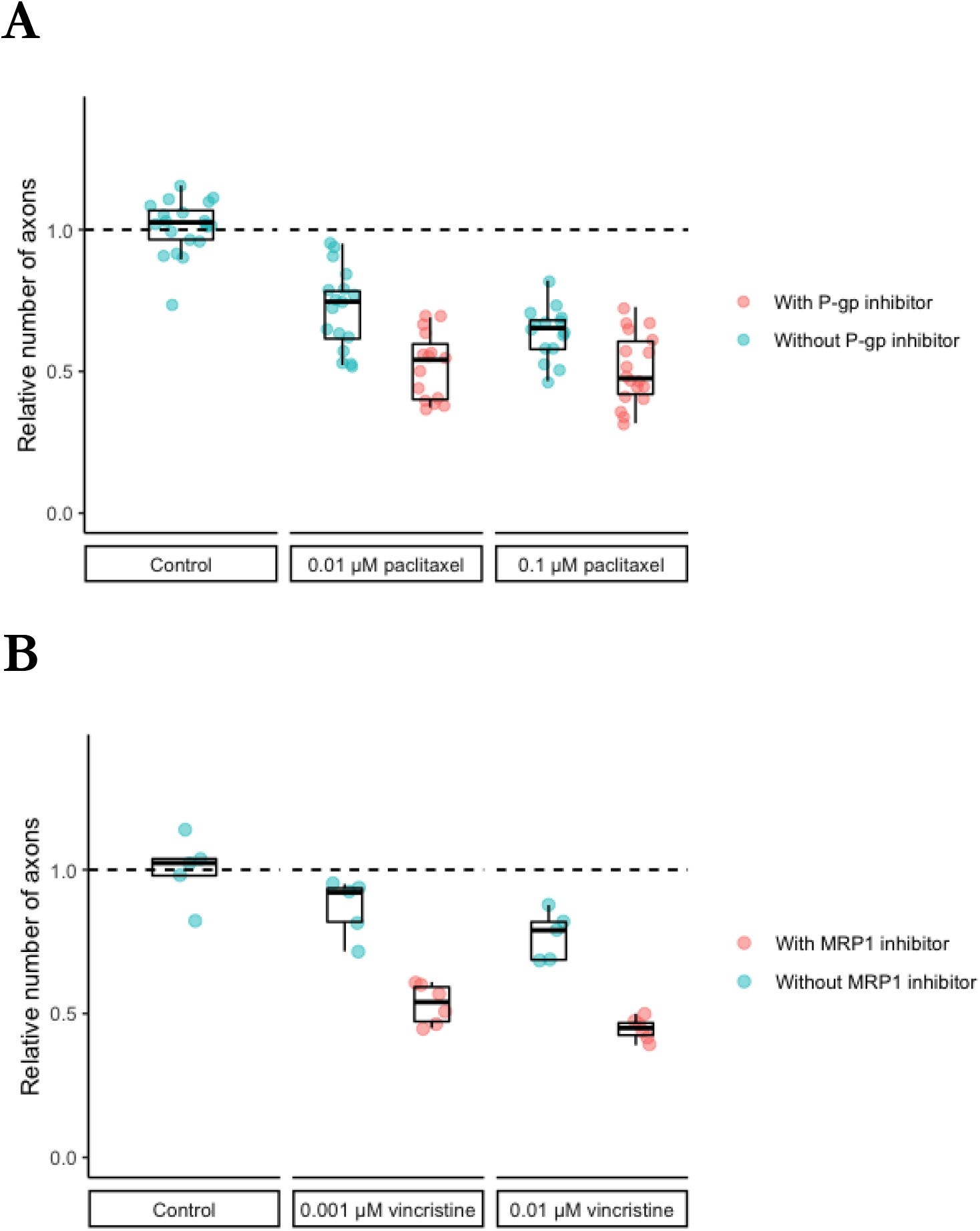
Inhibition of P-glycoprotein (P-gp) and multidrug assistance-associated protein 1 (MRP1) exacerbated paclitaxel and vincristine neurotoxicity in iPSC-SNs. Cells were pre-exposed to 4 µM valspodar or 4 µM MK-571 for 1 hour followed by 48 hours exposure with indicated concentrations of paclitaxel or vincristine. Cells were immunolabeled with peripherin, and the number of axons was quantified using Sholl analysis. Results are presented relative to vehicle control (n=1). Abbreviations: DMSO, dimethyl sulfoxide; iPSC-SNs, induced pluripotent stem cell-derived sensory neurons; MRP1, multidrug resistance-associated protein 1; P-gp, P-glycoprotein.

### Induction of P-gp as neuroprotection during neurotoxic chemotherapy

Our findings indicate that rifampicin induces *ABCB1* mRNA but not *ABCC1* mRNA in iPSC-SNs (**Figure S9**). To assess if the induction of *ABCB1* by rifampicin had a neuroprotective effect on neurotoxicity, 48 hours of induction was followed by 48 hours of exposure to paclitaxel or vincristine. Here, we found that pre-exposure to rifampicin preserved neuronal network integrity compared to paclitaxel or vincristine exposure alone (**Figures S10 and S11**). Consistent with this observation, rifampicin pre-exposure increased the number of axons and the neurite length in iPSC-SNs (**Figures 10 and S12**). Rifampicin pre-exposure protected against chemotherapy-induced transcriptional changes in iPSC-SN, as measured by the damage marker *ATF3* mRNA, and pain signaling as indicated by *TRPV1* mRNA (**Figure S13**).

**Figure 10.**
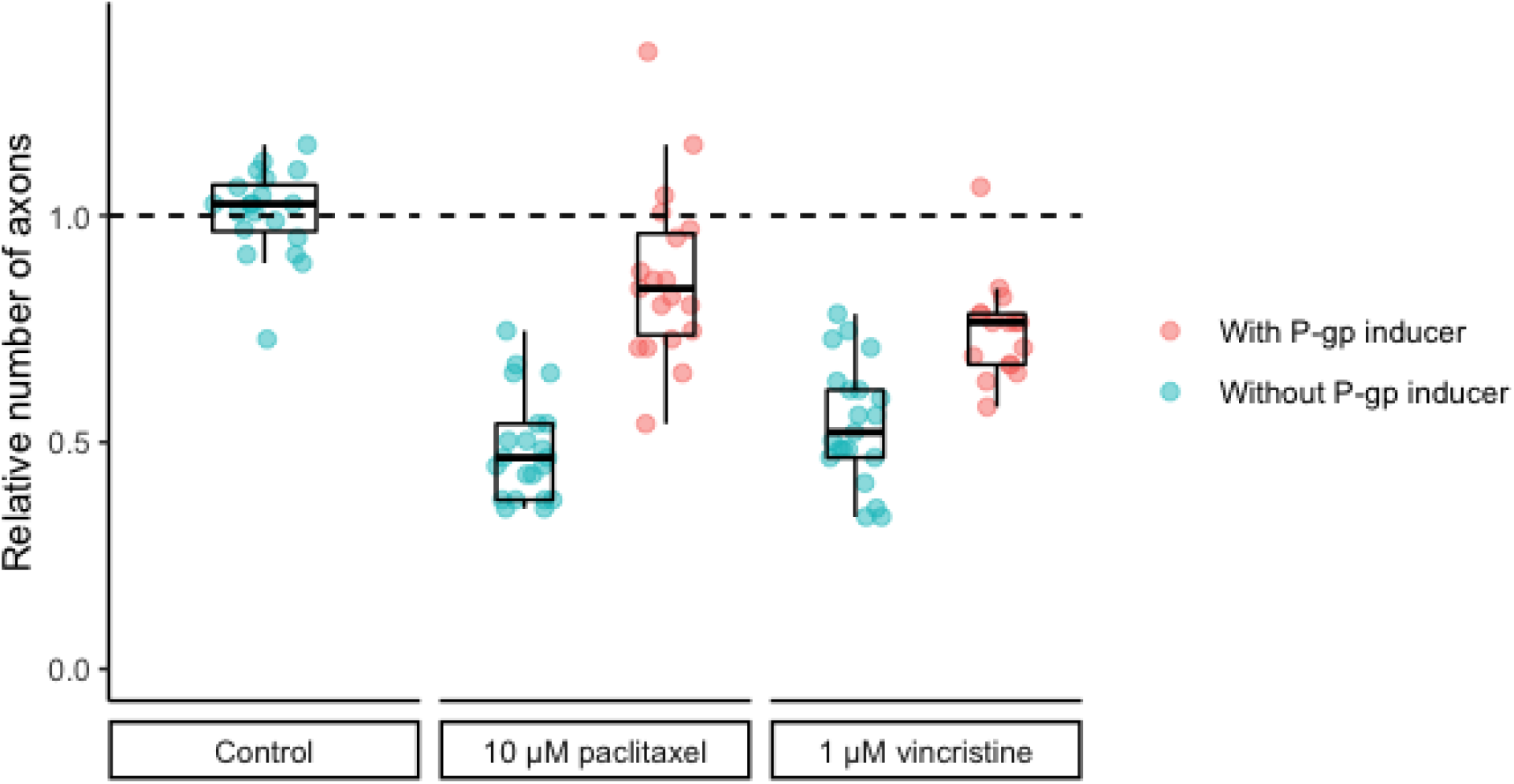
Rifampicin pre-exposure preserved neuronal network integrity of iPSC-SNs, as measured by the number of axons. iPSC-SNs were pre-exposed to rifampicin for 48 hours followed by 48 hours of exposure to the indicated concentrations of paclitaxel or vincristine. Cells were immunolabeled with peripherin, and the number of axons was quantified by Sholl analysis. Results are presented relative to vehicle control (n=1). Abbreviations: DMSO, dimethyl sulfoxide; iPSC-SNs, induced pluripotent stem cell-derived sensory neurons; P-gp, P-glycoprotein.

## Discussion

Paclitaxel and vincristine are among the most widely used chemotherapeutic agents to treat multiple types of cancer, such as breast cancer, lung cancer, leukemia, and lymphoma. The main dose-limiting adverse effect of both agents is CIPN, primarily due to lack of effective therapies for treatment and prevention. To provide insight into the molecular mechanisms underlying CIPN, we used a model of human sensory neurons derived from iPSCs. We confirmed that iPSC-SNs had a characteristic DRG morphology with ganglia-like structures and expressed canonical DRG markers. Paclitaxel and vincristine caused concentration-dependent neurotoxicity in iPSC-SNs. Both agents increased the mRNA expression of the pain receptor, *TRPV1*, and induced neuronal damage as measured by *ATF3* mRNA. Furthermore, we found that inhibition of efflux transporters exacerbated neurotoxicity in iPSC-SNs while induction of P-glycoprotein preserved neuronal network integrity and prevented transcriptional changes compared to neurotoxic chemotherapy alone.

Paclitaxel and vincristine caused distinct effects on the neuronal network which is in line with previous studies (15,24,31,32). The microtubule stabilization by paclitaxel leads to retraction and thickening of axons whereas vincristine destabilizes microtubules, leading to formation of microtubule fragments. These morphological effects are therefore specific to their distinct mechanism of action, though occurring without cell death (24,33–36). The effect of chemotherapy in iPSC-SNs might therefore reflect Wallerian degeneration where axons degenerate without affecting the cell bodies.

Our *in vitro* findings are consistent with skin biopsies from patients with chemotherapy-induced, diabetic, or hereditary neuropathy where the degeneration shows as loss of intraepidermal nerve fibers (IENFs), typically described as a “dying back” where axons degenerate in a distal-to-proximal manner (37,38). Consistent with previous findings in mice DRG neurons (15), we show that paclitaxel caused axonal retraction in iPSC-SNs that clearly reflected “dying back” degeneration. In contrast, the morphology of iPSC-SNs following vincristine exposure suggests fragmentation of both proximal and distal axons. Vincristine has been postulated to induce “dying back” degeneration, however, a previous study evaluating rat saphenous nerves found no difference between loss of proximal and distal axons (39). Another study showed that the number of axons in sural nerve biopsies from vincristine-treated mice was significantly lower distally (40). The observed degeneration in our *in vitro* model corresponds to the degeneration of IENFs in patient biopsies (41,42) and rodent models (43,44) treated with paclitaxel or vincristine. The loss of IENFs has also shown to be most pronounced in symptomatic areas of the skin following bortezomib treatment in cancer patients (45). The loss of IENFs could potentially be a major contributor to the development and persistence of CIPN.

TRPV1 is a well-known receptor that detects noxious stimuli in sensory neurons, leading to pain perception [41]. In the present study, we verified that TRPV1 was ubiquitously expressed in ganglia-like structures of iPSC-SNs and the exposure to paclitaxel and vincristine upregulated the mRNA expression of *TRPV1*. The functional properties of TRPV1 are known to be regulated by post-translational modifications e.g., phosphorylation leads to an upregulation of TRPV1 in paclitaxel-treated mice (46,47). It has previously been suggested that phosphorylation of TRPV1 can reduce the threshold for a stimulus to be detected as noxious (48). This might explain why paclitaxel and vincristine cause cancer patients to experience hypersensitivity to thermal and mechanical stimuli and hence, pain symptoms.

The knowledge regarding the role of efflux transporters in the distribution of chemotherapy in the PNS is scarce. P-gp and MRP1 are expressed in human DRG tissue (20) and corresponds to protein levels measured in our iPSC-SNs. Our findings agree with our recent study showing that P-gp inhibition increased intraneuronal accumulation and exacerbated neurotoxicity of paclitaxel in SH-SY5Y-derived neurons (20). We also showed that breast and ovarian cancer patients treated with P-gp inhibitors concomitantly with paclitaxel had a higher risk of dose modifications due to paclitaxel-induced peripheral neuropathy (20). These data are also supported by a clinical study showing that patients who were treated with a P-gp inhibitor during paclitaxel treatment had a higher risk of developing severe PIPN symptoms (49). The role of P-gp has previously been studied in rodents, though findings are conflicting perhaps due to species differences or other unknown factors (50–52). Both targeted and genome-wide association studies have linked *ABCB1* single nucleotide polymorphisms (SNPs) to risk of paclitaxel-induced peripheral neuropathy (53–55) and docetaxel-induced peripheral neuropathy (55). Variants in *ABCB1, ABCC1, and ABCC2* have also previously been linked to vincristine neurotoxicity (56–60). Here, we show that MRP1 inhibition leads to increased vincristine-induced neurotoxicity in iPSC-SNs. Our findings may provide a biological rationale for the correlation between *ABCC1* variants and vincristine-induced neurotoxicity. Collectively, P-gp and MRP1 appear to be critical for modulating sensory neuronal concentrations of chemotherapy and thus neurotoxicity.

To our knowledge, the induction of *ABCB1* has not been studied in the PNS previously. Rifampicin is the most potent inducer of P-gp and cytochrome P450 3A4 and affects both hepatic and intestinal drug metabolism (61). Here, we show that rifampicin induced *ABCB1* mRNA levels by 20% in iPSC-SNs which may confer increased efflux capacity, lower intraneuronal chemotherapy accumulation, and reduced neurotoxicity. Clinical studies are required to assess if the neuroprotection observed *in vitro* can be translated into clinical benefit. Importantly, local delivery to sensory neurons is required to avoid systemic exposure and protection of tumor cells.

The main limitation of this study is the lack of replications for inhibition and induction experiments and thus, the data should be interpreted with caution. Additionally, the experiments were performed with a single iPSC donor due to the comprehensive nature of the iPSC-SN differentiation. The clinical background of the donor is unknown and thus, future *in vitro* studies should utilize donors known to be tolerant or susceptible to neurotoxic chemotherapy. Another limitation is that iPSC-SNs were not co-cultured with Schwann cells that may influence chemotherapy-induced neurotoxicity through secretion of pro-inflammatory cytokines, and other factors (62).

The primary strength of this study is the use iPSC-SNs as a model representative of the DRG. iPSC-SNs contains various sensory neuron subtypes and recapitulate the DRG morphologically, transcriptionally, and functionally (24,63,64). Additionally, iPSC-SNs can overcome species differences and might improve translation of findings to clinical relevance compared to previously used models.

In conclusion, iPSC-SNs showed concentration-dependent neurotoxicity following exposure to paclitaxel and vincristine. The sensory neuron phenotypes were characterized by distinct morphological and transcriptional alterations, indicating their different mechanisms of action. Additionally, iPSC-SNs express the efflux transporters, P-gp and MRP1 and inhibition of these might exacerbate paclitaxel and vincristine neurotoxicity, respectively. We show that induction of P-gp by rifampicin might be a potential strategy to prevent chemotherapy-induced neurotoxicity. Although we identified P-gp as a potential therapeutic target, future studies can utilize iPSC-SNs in combination with hypothesis-free approaches, such as single-cell sequencing and CRISPR interference screen, to identify additional other therapeutic targets to meet the need for therapies for CIPN.

## Acknowledgements

We would like to thank Rasmus Andersen and Birgitte Damby Sørensen for their excellent analytical work.

## Author contribution

C.M. and T.B.S. designed the study. C.M. performed in vitro experiments. C.M., K.C.C., H.S.H., F.N., and O.P. analyzed the data, Å.F.S., D.L.K., and T.B.S. provided support and supervision. C.M., K.C.C., H.S.H., F.N., and T.B.S. wrote the manuscript. All authors reviewed the manuscript.

## Data availability

The data and R code are available on reasonable request from the corresponding author.

## References

1. Miller KD, Nogueira L, Devasia T, Mariotto AB, Yabroff KR, Jemal A, et al. Cancer treatment and survivorship statistics, 2022. CA Cancer J Clin. 2022 Sep;72(5):409–36.

2. Seretny M, Currie GL, Sena ES, Ramnarine S, Grant R, MacLeod MR, et al. Incidence, prevalence, and predictors of chemotherapy-induced peripheral neuropathy: A systematic review and meta-analysis. Pain. 2014 Dec;155(12):2461–70.

3. Mortensen C, Andersen NE, Stage TB. Bridging the Translational Gap in Chemotherapy-Induced Peripheral Neuropathy with iPSC-Based Modeling. Cancers (Basel). 2022 Aug 15;14(16):3939.

4. Fukuda Y, Li Y, Segal RA. A Mechanistic Understanding of Axon Degeneration in Chemotherapy-Induced Peripheral Neuropathy. Front Neurosci. 2017;11:481.

5. Kober KM, Mazor M, Abrams G, Olshen A, Conley YP, Hammer M, et al. Phenotypic Characterization of Paclitaxel-Induced Peripheral Neuropathy in Cancer Survivors. J Pain Symptom Manage. 2018 Dec;56(6):908-919.e3.

6. Mizrahi D, Park SB, Li T, Timmins HC, Trinh T, Au K, et al. Hemoglobin, Body Mass Index, and Age as Risk Factors for Paclitaxel- and Oxaliplatin-Induced Peripheral Neuropathy. JAMA Network Open. 2021 Feb 15;4(2):e2036695.

7. Cavaletti G, Alberti P, Frigeni B, Piatti M, Susani E. Chemotherapy-induced neuropathy. Curr Treat Options Neurol. 2011 Apr;13(2):180–90.

8. Loprinzi CL, Lacchetti C, Bleeker J, Cavaletti G, Chauhan C, Hertz DL, et al. Prevention and Management of Chemotherapy-Induced Peripheral Neuropathy in Survivors of Adult Cancers: ASCO Guideline Update. J Clin Oncol. 2020 Oct 1;38(28):3325–48.

9. Gornstein E, Schwarz TL. The paradox of paclitaxel neurotoxicity: Mechanisms and unanswered questions. Neuropharmacology. 2014 Jan 1;76:175–83.

10. LaPointe NE, Morfini G, Brady ST, Feinstein SC, Wilson L, Jordan MA. Effects of eribulin, vincristine, paclitaxel and ixabepilone on fast axonal transport and kinesin-1 driven microtubule gliding: Implications for chemotherapy-induced peripheral neuropathy. NeuroToxicology. 2013 Jul 1;37:231–9.

11. Zheng H, Xiao WH, Bennett GJ. Functional deficits in peripheral nerve mitochondria in rats with paclitaxel- and oxaliplatin-evoked painful peripheral neuropathy. Experimental Neurology. 2011 Dec 1;232(2):154–61.

12. Makker PGS, Duffy SS, Lees JG, Perera CJ, Tonkin RS, Butovsky O, et al. Characterisation of Immune and Neuroinflammatory Changes Associated with Chemotherapy-Induced Peripheral Neuropathy. PLoS One. 2017 Jan 26;12(1):e0170814.

13. Zhang H, Dougherty PM. Enhanced Excitability of Primary Sensory Neurons and Altered Gene Expression of Neuronal Ion Channels in Dorsal Root Ganglion in Paclitaxel-Induced Peripheral Neuropathy. Anesthesiology. 2014 Jun;120(6):1463–75.

14. Park SB, Goldstein D, Krishnan AV, Lin CSY, Friedlander ML, Cassidy J, et al. Chemotherapy-induced peripheral neurotoxicity: a critical analysis. CA Cancer J Clin. 2013 Dec;63(6):419–37.

15. Gornstein EL, Schwarz TL. Neurotoxic mechanisms of paclitaxel are local to the distal axon and independent of transport defects. Exp Neurol. 2017 Feb;288:153–66.

16. Silva A, Wang Q, Wang M, Ravula SK, Glass JD. Evidence for direct axonal toxicity in vincristine neuropathy. J Peripher Nerv Syst. 2006 Sep;11(3):211–6.

17. Holzer AK, Karreman C, Suciu I, Furmanowsky LS, Wohlfarth H, Loser D, et al. Generation of human nociceptor-enriched sensory neurons for the study of pain-related dysfunctions [Internet]. bioRxiv; 2022 [cited 2022 May 18]. p. 2022.02.19.480828. Available from: https://www.biorxiv.org/content/10.1101/2022.02.19.480828v1

18. Chan A, Hertz DL, Morales M, Adams EJ, Gordon S, Tan CJ, et al. Biological Predictors of Chemotherapy Induced Peripheral Neuropathy (CIPN): MASCC Neurological Complications Working Group Overview. Support Care Cancer. 2019 Oct;27(10):3729–37.

19. Löscher W, Potschka H. Blood-Brain Barrier Active Efflux Transporters: ATP-Binding Cassette Gene Family. NeuroRx. 2005 Jan;2(1):86–98.

20. Stage TB, Mortensen C, Khalaf S, Steffensen V, Hammer HS, Xiong C, et al. P-Glycoprotein Inhibition Exacerbates Paclitaxel Neurotoxicity in Neurons and Patients With Cancer. Clin Pharmacol Ther. 2020 Sep;108(3):671–80.

21. Agergaard K, Mau-Sørensen M, Stage TB, Jørgensen TL, Hassel RE, Steffensen KD, et al. Clopidogrel-Paclitaxel Drug-Drug Interaction: A Pharmacoepidemiologic Study. Clin Pharmacol Ther. 2017 Sep;102(3):547–53.

22. Mealey KL, Fidel J. P‐Glycoprotein Mediated Drug Interactions in Animals and Humans with Cancer. J Vet Intern Med. 2015;29(1):1–6.

23. Mortensen C, Steffensen KD, Simonsen E, Herskind K, Madsen JS, Olsen DA, et al. Neurofilament light chain as a biomarker of axonal damage in sensory neurons and paclitaxel-induced peripheral neuropathy in ovarian cancer patients. Pain. 2022 Dec 9;

24. Xiong C, Chua KC, Stage TB, Priotti J, Kim J, Altman-Merino A, et al. Human Induced Pluripotent Stem Cell Derived Sensory Neurons are Sensitive to the Neurotoxic Effects of Paclitaxel. Clin Transl Sci. 2020 Dec 19;

25. Chua KC, Xiong C, Ho C, Mushiroda T, Jiang C, Mulkey F, et al. Genomewide Meta-Analysis Validates a Role for S1PR1 in Microtubule Targeting Agent-Induced Sensory Peripheral Neuropathy. Clinical Pharmacology & Therapeutics. 2020;108(3):625–34.

26. Schmittgen TD, Livak KJ. Analyzing real-time PCR data by the comparative CT method. Nat Protoc. 2008 Jun;3(6):1101–8.

27. Weiß F, Hammer HS, Klein K, Planatscher H, Zanger UM, Norén A, et al. Direct Quantification of Cytochromes P450 and Drug Transporters-A Rapid, Targeted Mass Spectrometry-Based Immunoassay Panel for Tissues and Cell Culture Lysates. Drug Metab Dispos. 2018 Apr;46(4):387–96.

28. Belghit I, Varunjikar M, Lecrenier MC, Steinhilber A, Niedzwiecka A, Wang YV, et al. Future feed control – Tracing banned bovine material in insect meal. Food Control. 2021 Oct 1;128:108183.

29. Wuerger LTD, Hammer HS, Hofmann U, Kudiabor F, Sieg H, Braeuning A. Okadaic acid influences xenobiotic metabolism in HepaRG cells. EXCLI J. 2022 Aug 1;21:1053–65.

30. Seijffers R, Mills CD, Woolf CJ. ATF3 increases the intrinsic growth state of DRG neurons to enhance peripheral nerve regeneration. J Neurosci. 2007 Jul 25;27(30):7911–20.

31. Geisler S, Doan RA, Cheng GC, Cetinkaya-Fisgin A, Huang SX, Höke A, et al. Vincristine and bortezomib use distinct upstream mechanisms to activate a common SARM1-dependent axon degeneration program. JCI Insight. 2019 Sep 5;4(17):e129920, 129920.

32. Akin EJ, Alsaloum M, Higerd GP, Liu S, Zhao P, Dib-Hajj FB, et al. Paclitaxel increases axonal localization and vesicular trafficking of Nav1.7. Brain. 2021 Jul 28;144(6):1727–37.

33. Vojnits K, Mahammad S, Collins TJ, Bhatia M. Chemotherapy-Induced Neuropathy and Drug Discovery Platform Using Human Sensory Neurons Converted Directly from Adult Peripheral Blood. Stem Cells Transl Med. 2019 Nov;8(11):1180–91.

34. Rana P, Luerman G, Hess D, Rubitski E, Adkins K, Somps C. Utilization of iPSC-derived human neurons for high-throughput drug-induced peripheral neuropathy screening. Toxicol In Vitro. 2017 Dec;45(Pt 1):111–8.

35. Wing C, Komatsu M, Delaney SM, Krause M, Wheeler HE, Dolan ME. Application of stem cell derived neuronal cells to evaluate neurotoxic chemotherapy. Stem Cell Res. 2017 Jul;22:79–88.

36. Wang M, Wang J, Tsui AYP, Li Z, Zhang Y, Zhao Q, et al. Mechanisms of peripheral neurotoxicity associated with four chemotherapy drugs using human induced pluripotent stem cell-derived peripheral neurons. Toxicol In Vitro. 2021 Dec;77:105233.

37. Hartmannsberger B, Doppler K, Stauber J, Schlotter-Weigel B, Young P, Sereda MW, et al. Intraepidermal nerve fibre density as biomarker in Charcot–Marie–Tooth disease type 1A. Brain Communications. 2020 Jan 1;2(1):fcaa012.

38. Beiswenger KK, Calcutt NA, Mizisin AP. Epidermal Nerve Fiber Quantification in the Assessment of Diabetic Neuropathy. Acta Histochem. 2008;110(5):351–62.

39. Cottschalk PG, Dyck PJ, Kiely JM. Vinca alkaloid neuropathy: Nerve biopsy studies in rats and in man. Neurology. 1968 Sep 1;18(9):875–875.

40. Geisler S, Doan RA, Strickland A, Huang X, Milbrandt J, DiAntonio A. Prevention of vincristine-induced peripheral neuropathy by genetic deletion of SARM1 in mice. Brain. 2016 Dec;139(Pt 12):3092–108.

41. Boyette-Davis JA, Cata JP, Driver LC, Novy DM, Bruel BM, Mooring DL, et al. Persistent chemoneuropathy in patients receiving the plant alkaloids paclitaxel and vincristine. Cancer Chemother Pharmacol. 2013 Mar;71(3):619–26.

42. Sahenk Z, Barohn R, New P, Mendell JR. Taxol neuropathy. Electrodiagnostic and sural nerve biopsy findings. Arch Neurol. 1994 Jul;51(7):726–9.

43. Boyette-Davis J, Xin W, Zhang H, Dougherty PM. Intraepidermal nerve fiber loss corresponds to the development of taxol-induced hyperalgesia and can be prevented by treatment with minocycline. Pain. 2011 Feb;152(2):308–13.

44. Siau C, Xiao W, Bennett GJ. Paclitaxel- and vincristine-evoked painful peripheral neuropathies: loss of epidermal innervation and activation of Langerhans cells. Exp Neurol. 2006 Oct;201(2):507–14.

45. Boyette-Davis JA, Cata JP, Zhang H, Driver LC, Wendelschafer-Crabb G, Kennedy WR, et al. Follow-up psychophysical studies in bortezomib-related chemoneuropathy patients. J Pain. 2011 Sep;12(9):1017–24.

46. Bhave G, Zhu W, Wang H, Brasier DJ, Oxford GS, Gereau RW. cAMP-dependent protein kinase regulates desensitization of the capsaicin receptor (VR1) by direct phosphorylation. Neuron. 2002 Aug 15;35(4):721–31.

47. Zhuang ZY, Xu H, Clapham DE, Ji RR. Phosphatidylinositol 3-kinase activates ERK in primary sensory neurons and mediates inflammatory heat hyperalgesia through TRPV1 sensitization. J Neurosci. 2004 Sep 22;24(38):8300–9.

48. Palazzo E, Luongo L, de Novellis V, Rossi F, Marabese I, Maione S. Transient receptor potential vanilloid type 1 and pain development. Curr Opin Pharmacol. 2012 Feb;12(1):9–17.

49. Lhommé C, Joly F, Walker JL, Lissoni AA, Nicoletto MO, Manikhas GM, et al. Phase III study of valspodar (PSC 833) combined with paclitaxel and carboplatin compared with paclitaxel and carboplatin alone in patients with stage IV or suboptimally debulked stage III epithelial ovarian cancer or primary peritoneal cancer. J Clin Oncol. 2008 Jun 1;26(16):2674–82.

50. Saito T, Zhang ZJ, Ohtsubo T, Noda I, Shibamori Y, Yamamoto T, et al. Homozygous disruption of the mdrla P-glycoprotein gene affects blood-nerve barrier function in mice administered with neurotoxic drugs. Acta Otolaryngol. 2001 Sep;121(6):735–42.

51. Huang L, Li X, Roberts J, Janosky B, Lin MHJ. Differential role of P-glycoprotein and breast cancer resistance protein in drug distribution into brain, CSF and peripheral nerve tissues in rats. Xenobiotica. 2015;45(6):547–55.

52. Liu H, Chen Y, Huang L, Sun X, Fu T, Wu S, et al. Drug Distribution into Peripheral Nerve. J Pharmacol Exp Ther. 2018 May;365(2):336–45.

53. Sissung TM, Mross K, Steinberg SM, Behringer D, Figg WD, Sparreboom A, et al. Association of ABCB1 genotypes with paclitaxel-mediated peripheral neuropathy and neutropenia. Eur J Cancer. 2006 Nov;42(17):2893–6.

54. Tanabe Y, Shimizu C, Hamada A, Hashimoto K, Ikeda K, Nishizawa D, et al. Paclitaxel-induced sensory peripheral neuropathy is associated with an ABCB1 single nucleotide polymorphism and older age in Japanese. Cancer Chemother Pharmacol. 2017 Jun;79(6):1179–86.

55. Kus T, Aktas G, Kalender ME, Demiryurek AT, Ulasli M, Oztuzcu S, et al. Polymorphism of CYP3A4 and ABCB1 genes increase the risk of neuropathy in breast cancer patients treated with paclitaxel and docetaxel. Onco Targets Ther. 2016 Aug 12;9:5073–80.

56. Wright GEB, Amstutz U, Drögemöller BI, Shih J, Rassekh SR, Hayden MR, et al. Pharmacogenomics of Vincristine-Induced Peripheral Neuropathy Implicates Pharmacokinetic and Inherited Neuropathy Genes. Clin Pharmacol Ther. 2019;105(2):402–10.

57. Lopez-Lopez E, Gutierrez-Camino A, Astigarraga I, Navajas A, Echebarria-Barona A, Garcia-Miguel P, et al. Vincristine pharmacokinetics pathway and neurotoxicity during early phases of treatment in pediatric acute lymphoblastic leukemia. Pharmacogenomics. 2016 May;17(7):731–41.

58. Ceppi F, Langlois-Pelletier C, Gagné V, Rousseau J, Ciolino C, Lorenzo SD, et al. Polymorphisms of the vincristine pathway and response to treatment in children with childhood acute lymphoblastic leukemia. Pharmacogenomics. 2014 Jun;15(8):1105–16.

59. Broyl A, Corthals SL, Jongen JL, van der Holt B, Kuiper R, de Knegt Y, et al. Mechanisms of peripheral neuropathy associated with bortezomib and vincristine in patients with newly diagnosed multiple myeloma: a prospective analysis of data from the HOVON-65/GMMG-HD4 trial. Lancet Oncol. 2010 Nov;11(11):1057–65.

60. Franca R, Rebora P, Bertorello N, Fagioli F, Conter V, Biondi A, et al. Pharmacogenetics and induction/consolidation therapy toxicities in acute lymphoblastic leukemia patients treated with AIEOP-BFM ALL 2000 protocol. Pharmacogenomics J. 2017 Jan;17(1):4–10.

61. Niemi M, Backman JT, Fromm MF, Neuvonen PJ, Kivistö KT. Pharmacokinetic interactions with rifampicin : clinical relevance. Clin Pharmacokinet. 2003;42(9):819–50.

62. Wang XM, Lehky TJ. Discovering Cytokines as Targets for Chemotherapy-Induced Painful Peripheral Neuropathy. Cytokine. 2012 Jul;59(1):3–9.

63. Schinke C, Vallone VF, Ivanov A, Peng Y, Körtvelyessy P, Nolte L, et al. Modeling chemotherapy induced neurotoxicity with human induced pluripotent stem cell (iPSC)-derived sensory neurons. Neurobiology of Disease. 2021 May 11;105391.

64. Schwartzentruber J, Foskolou S, Kilpinen H, Rodrigues J, Alasoo K, Knights AJ, et al. Molecular and functional variation in iPSC-derived sensory neurons. Nat Genet. 2018 Jan;50(1):54–61.

