## Supplementary materials for "Paclitaxel- and vincristine-induced neurotoxicity and drug transport in sensory neurons"

Mortensen *et al.* 2022

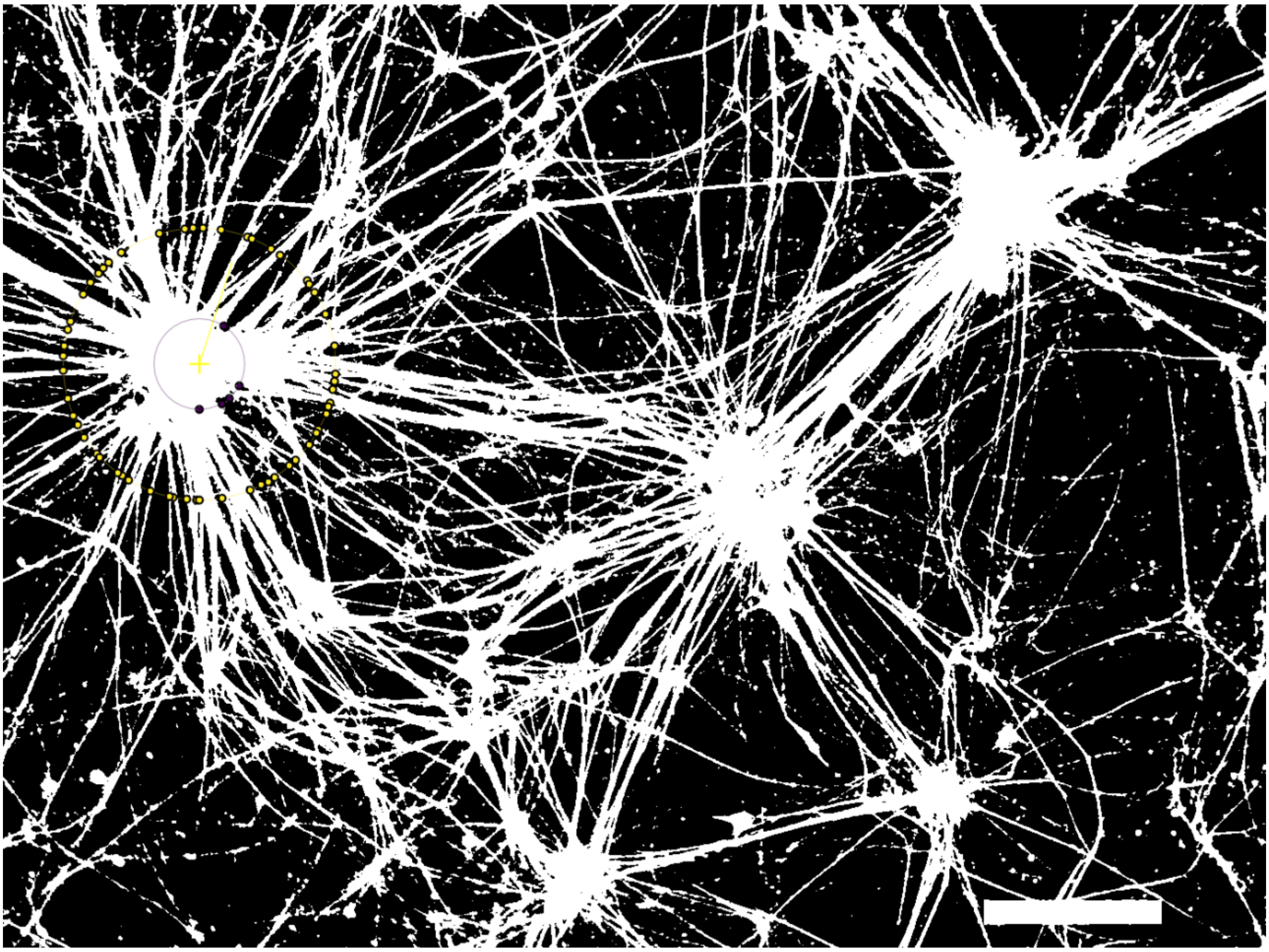

**Figure S1**. Threshold adjustment and Sholl analysis of a single ganglion. iPSC-derived sensory neurons were immunolabeled for peripherin, and Sholl analysis were performed in ImageJ using an appropriate threshold to visualize the neuronal network. The center of each ganglion was defined using the straight-line tool, and the number of axons was then quantified at an appropriate end radius.

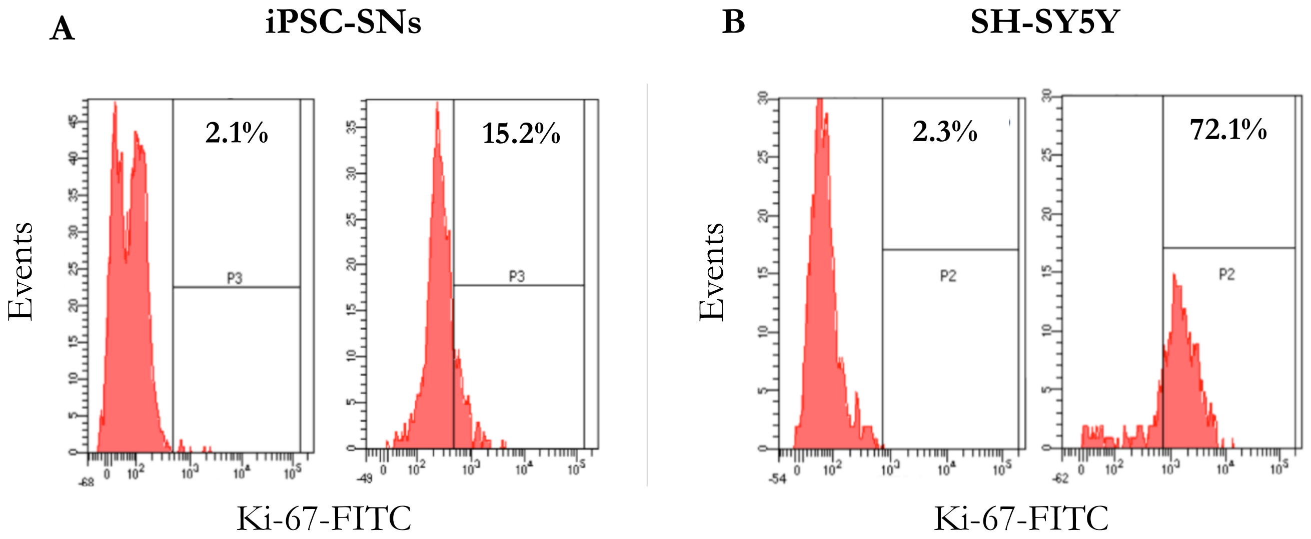


**Figure S2**. Flowcytometric analysis of iPSC-SNs on day 12 (A). Undifferentiated SH-SY5Y neuroblastoma cells was included as a positive control (B). The cell proliferation status was detected by the expression of Ki-67. Abbreviations: FITC, fluorescein isothiocyanate; iPSC-SNs, induced pluripotent stem cell-derived sensory neurons.

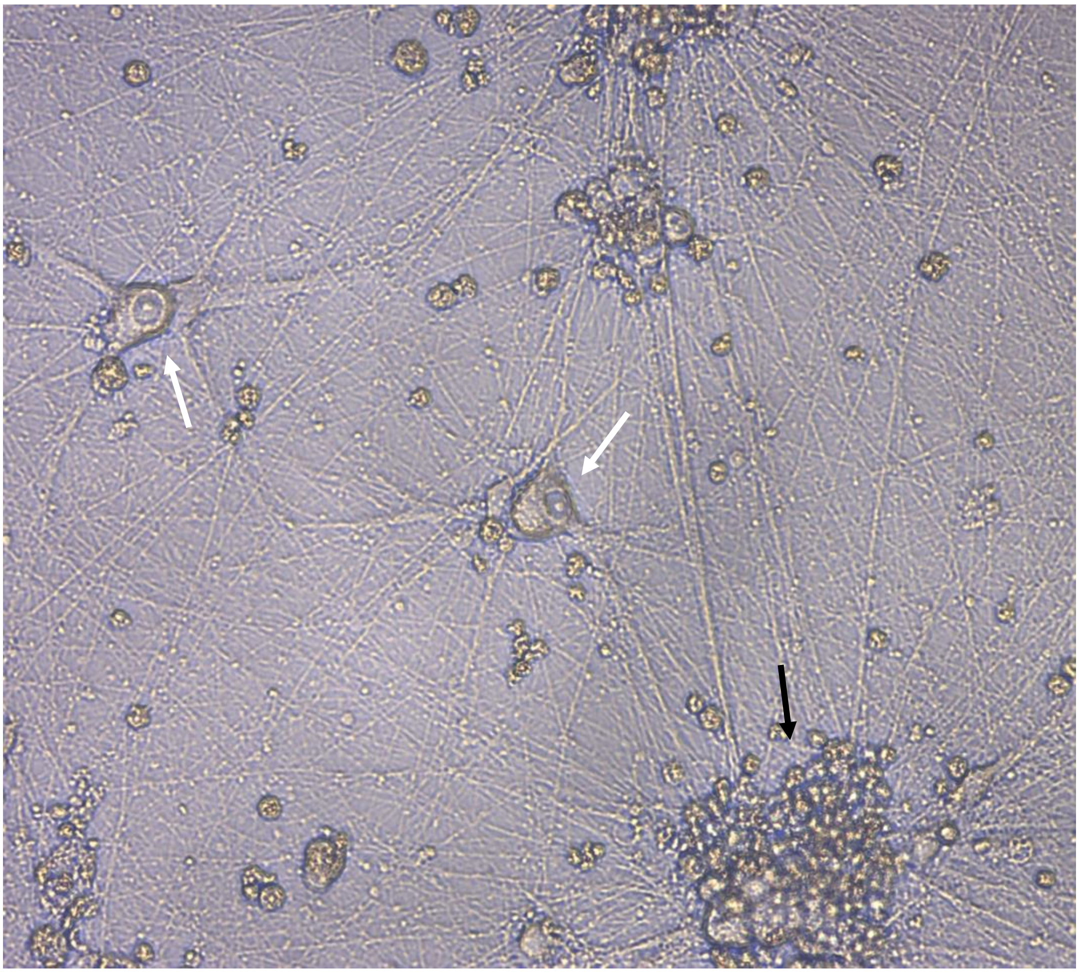

**Figure S3.** The cell bodies of iPSC-derived sensory neurons are mainly localized in large ganglia-like structures (black arrowhead), but single sensory neurons can also be observed in the culture (white arrowheads). Image was captured using VisiScope IT414 with the 10X objective.

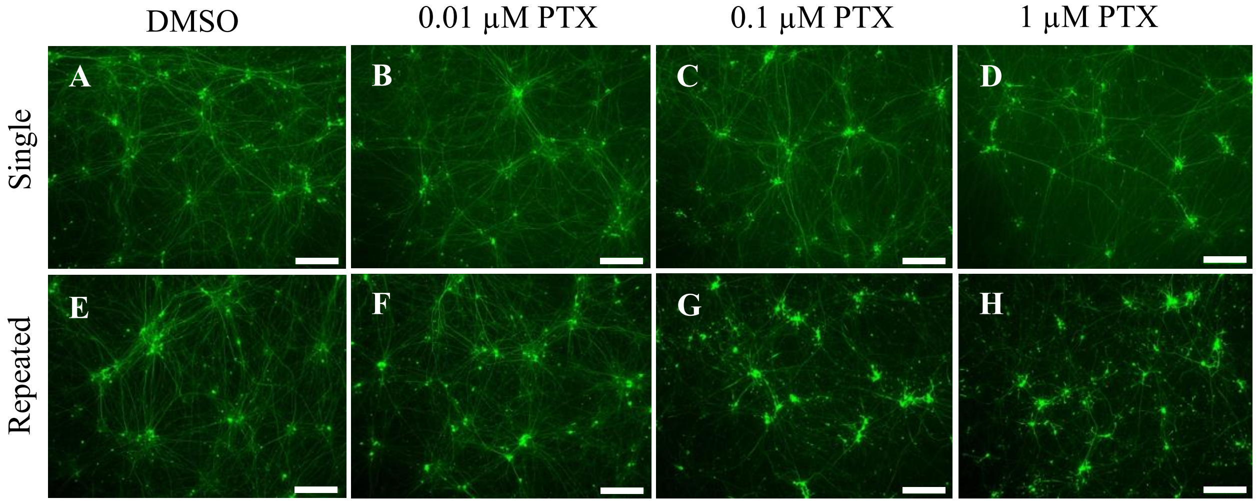


**Figure S4**. Repeated exposure of paclitaxel increases the axonal retraction and reduces the neuronal network of iPSC-SNs. For single exposure, cells were exposed to DMSO, 0.01 μM, 0.1 μM, and 1 μM paclitaxel for 48 hours (A-D). For repeated exposure, iPSC-SNs were treated in parallel for 48 hours followed by change to paclitaxel-free medium for 5 days and then cells were treated with paclitaxel for further 48 hours (E-H). Cells were fixed and labeled with peripherin. Images were captured using the Leica DMI 4000B microscope with a 10x objective. Scale bars represent 200 μm. Abbreviations: DMSO, dimethyl sulfoxide; iPSC-SNs, induced pluripotent stem cell-derived sensory neurons; PTX, paclitaxel.

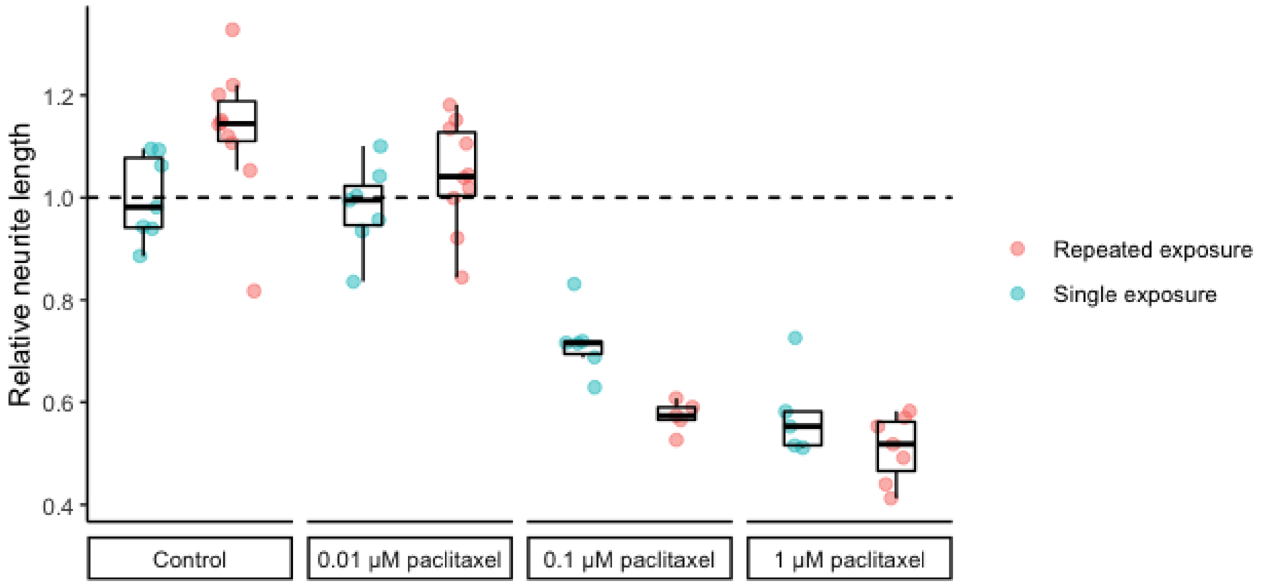


**Figure S5.** Repeated exposure of paclitaxel reduces the neurite length in iPSC-SNs**.** For single exposure, iPSC-SNs were treated with DMSO, 0.01 µM, 0.1 µM or 1 µM paclitaxel for 48 hours on day 38 of neuronal differentiation. For repeated exposure, iPSC-SNs were exposed to paclitaxel for 48 hours. After removal of the paclitaxel-containing medium, new N2 medium was added and after 5 days, cells were treated with paclitaxel for further 48 hours. iPSC-SNs were fixed and immunolabeled for peripherin to visualize the neuronal network. The immunolabeled images were analyzed in MIPAR. The results are shown relative to DMSO control (single exposure). Abbreviations: DMSO, dimethyl sulfoxide; iPSC-SNs, induced pluripotent stem cell-derived sensory neurons.

**A**

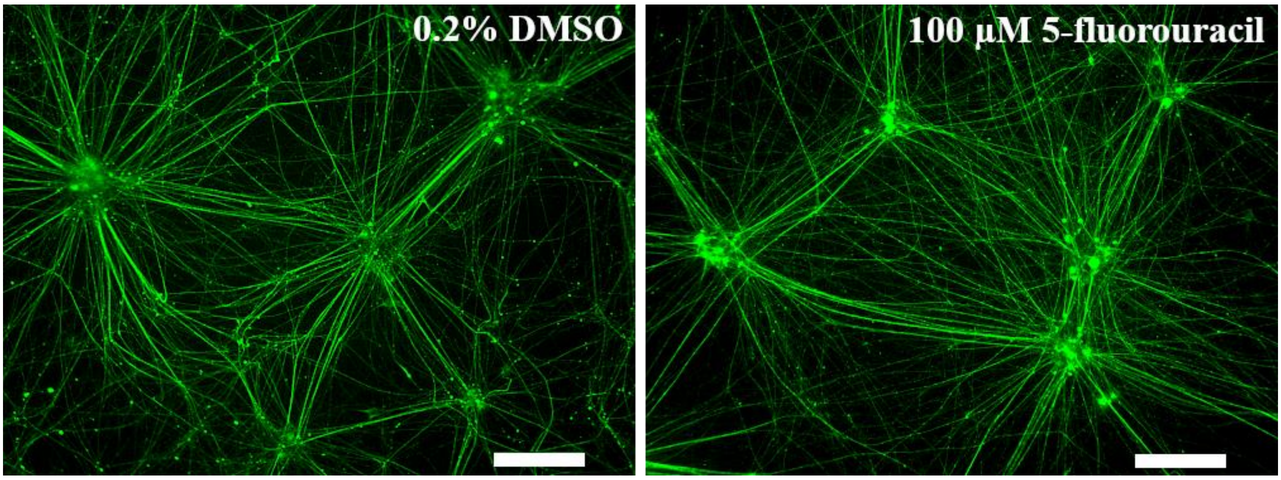


**B**

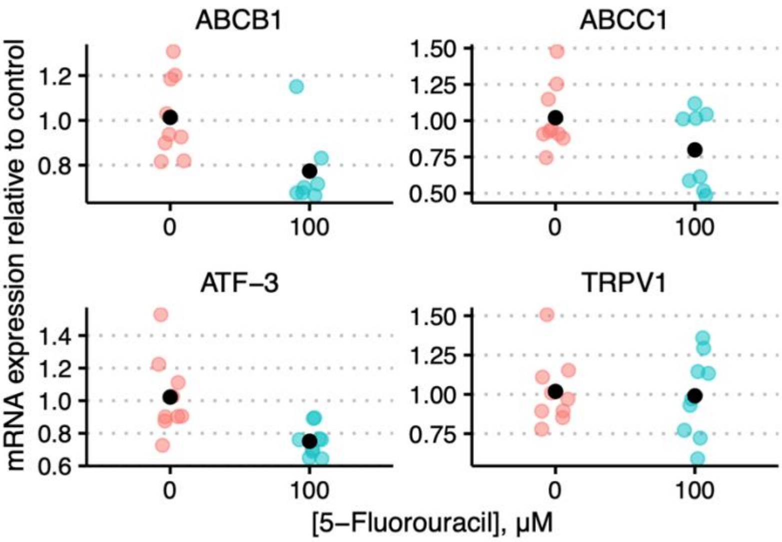


**Figure S6.** 5-flourouracil did not affect the neuronal network or *TRPV1* mRNA levels in iPSC-derived sensory neurons. The mRNA expression of *ATF3*, *ABCB1*, and *ABCC1* was reduced by 5-flourouracil. Mature iPSC-derived sensory neurons were treated with vehicle (0.2% DMSO) and 100 μM 5-fluorouracil for 48 hours followed by immunolabeling with peripherin (A). Images were captured using the Leica DMI 4000B microscope with the 10X objective. Scale bars represent 200 μm. The mRNA level is presented relative to vehicle from each differentiation (B). The black dot represents the mean value of 3 wells per condition from the 3 independent differentiations.

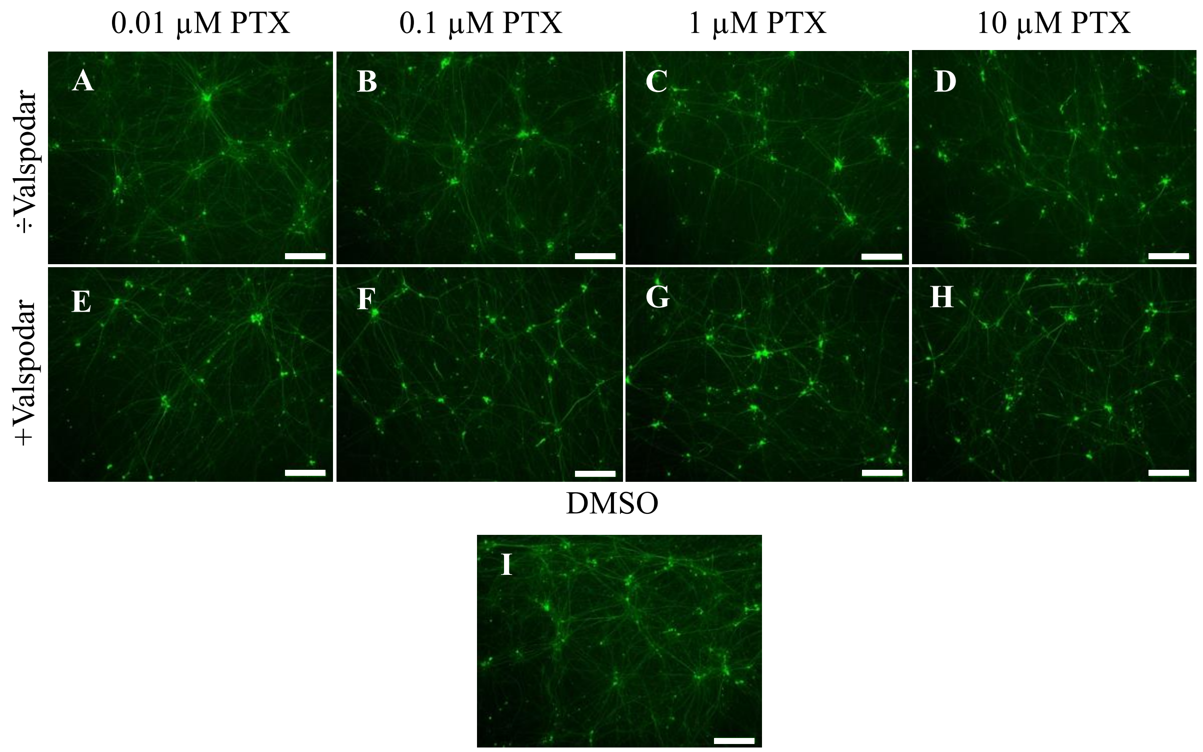


**Figure S7.** Inhibition of P-glycoprotein (P-gp) reduced the neuronal network of iPSC-SNs. Cells were exposed to vehicle (0.2% DMSO) and indicated concentrations of paclitaxel for 48 hours. For inhibition of P-gp, iPSC-SNs were pre-exposed to 4 μM valspodar for 1 hour followed by paclitaxel exposure for 48 hours. Scale bars represent 200 μm. Abbreviations: DMSO, dimethyl sulfoxide; iPSC-SNs, induced pluripotent stem cell-derived sensory neurons; PTX, paclitaxel.

A

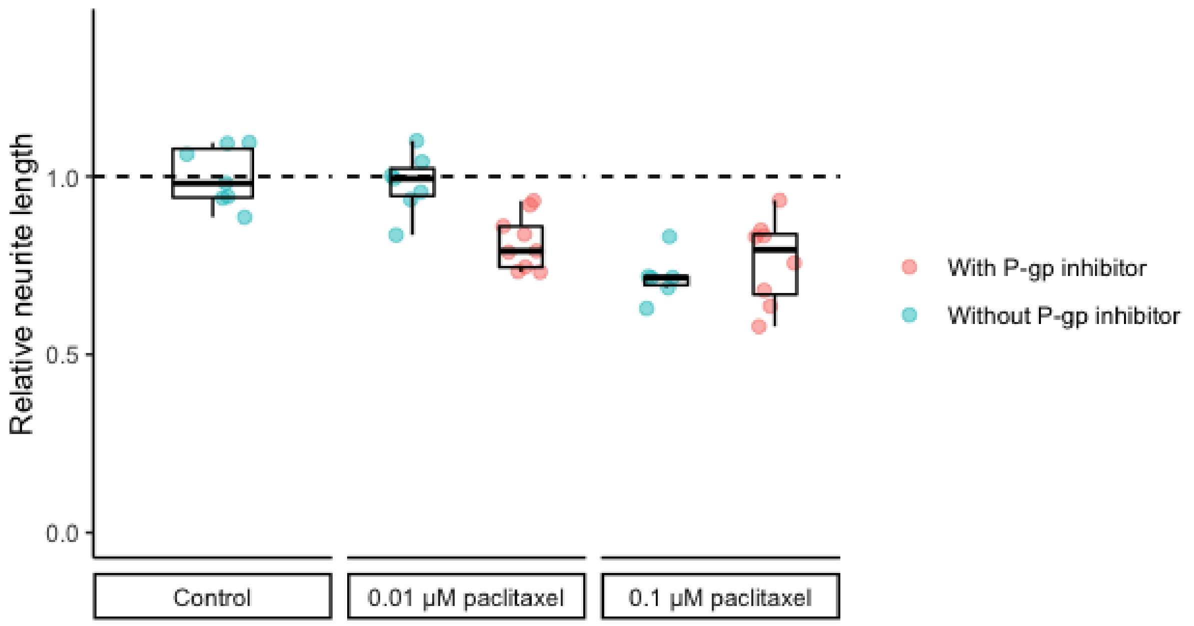


B

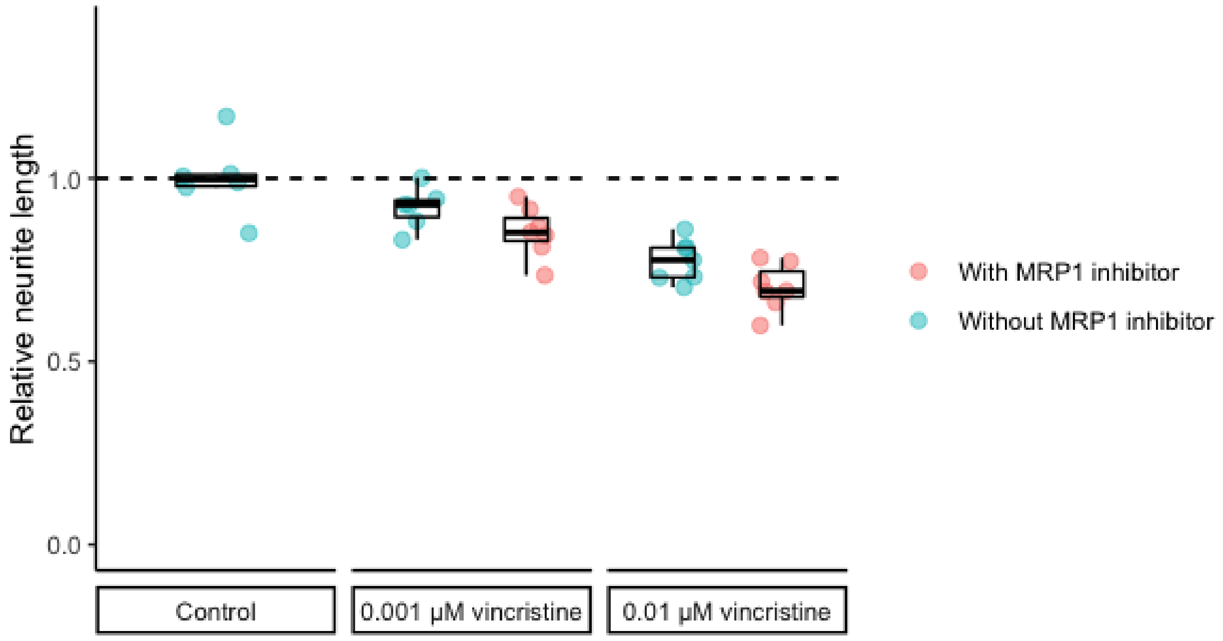


**Figure S8.** Inhibition of P-glycoprotein (P-gp) exacerbated paclitaxel neurotoxicity in iPSC-SNs (A). Inhibition of multidrug resistance-associated protein 1 (MRP1) exacerbated vincristine neurotoxicity in iPSC-SNs. (A) Cells were pretreated with 4 μM valspodar for 1 hours followed by 48 hours treatment with indicated concentrations of paclitaxel. (B) Cells were pretreated with 4 μM MK-571 for 1 hour followed by 48 hours treatment with indicated concentrations of vincristine. Cells were immunolabeled with peripherin and neurite length was quantified using MIPAR. Results are presented relative to DMSO control. Abbreviations: DMSO, dimethyl sulfoxide; iPSC-SNs, induced pluripotent stem cell-derived sensory neurons.

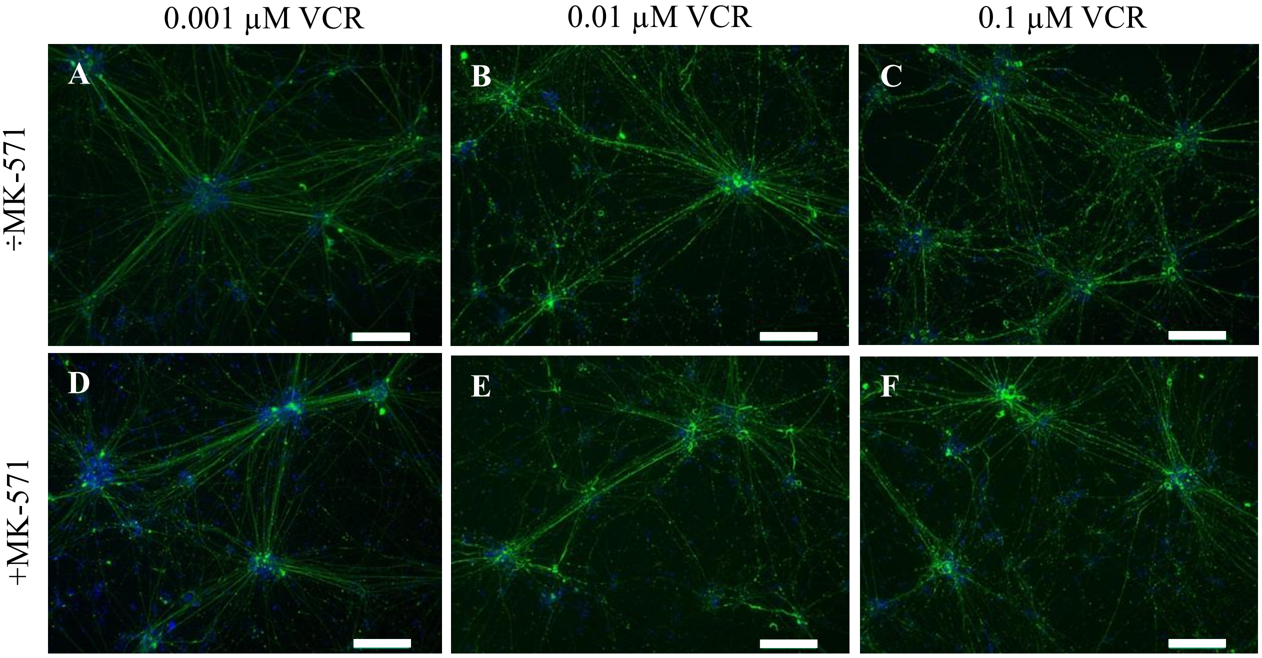


**Figure S9.** Inhibition of MRP1 exacerbated axonal fragmentation in iPSC-SNs. Cells were exposed to indicated concentrations of vincristine for 48 hours. For inhibition of MRP1, iPSC-SNs were pre-exposed to 4 μM MK-571 for 1 hour followed by vincristine exposure for 48 hours. Scale bars represent 200 μm. Abbreviations: iPSC-SNs, induced pluripotent stem cell-derived sensory neurons; VCR, vincristine.

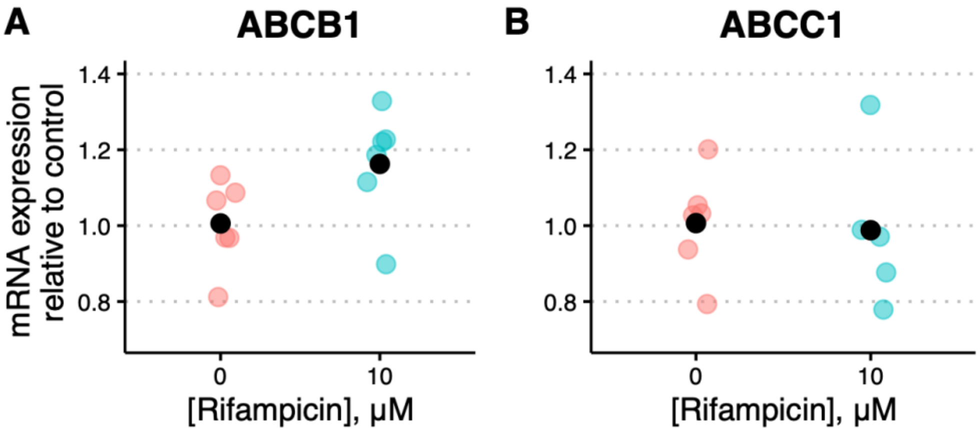


**Figure S10.** Rifampicin induces *ABCB1* but not *ABCC1* mRNA levels in iPSC-SNs. Cells were treated with vehicle (0.2% DMSO) and 10 µM rifampicin for 48 hours. The mRNA level is presented relative to vehicle from each differentiation. The black dot represents the mean value of 3 wells per condition from the 2 independent differentiations.

Abbreviations: ABC, ATP-binding cassette; DMSO, dimethyl sulfoxide, induced pluripotent stem cell-derived sensory neurons.

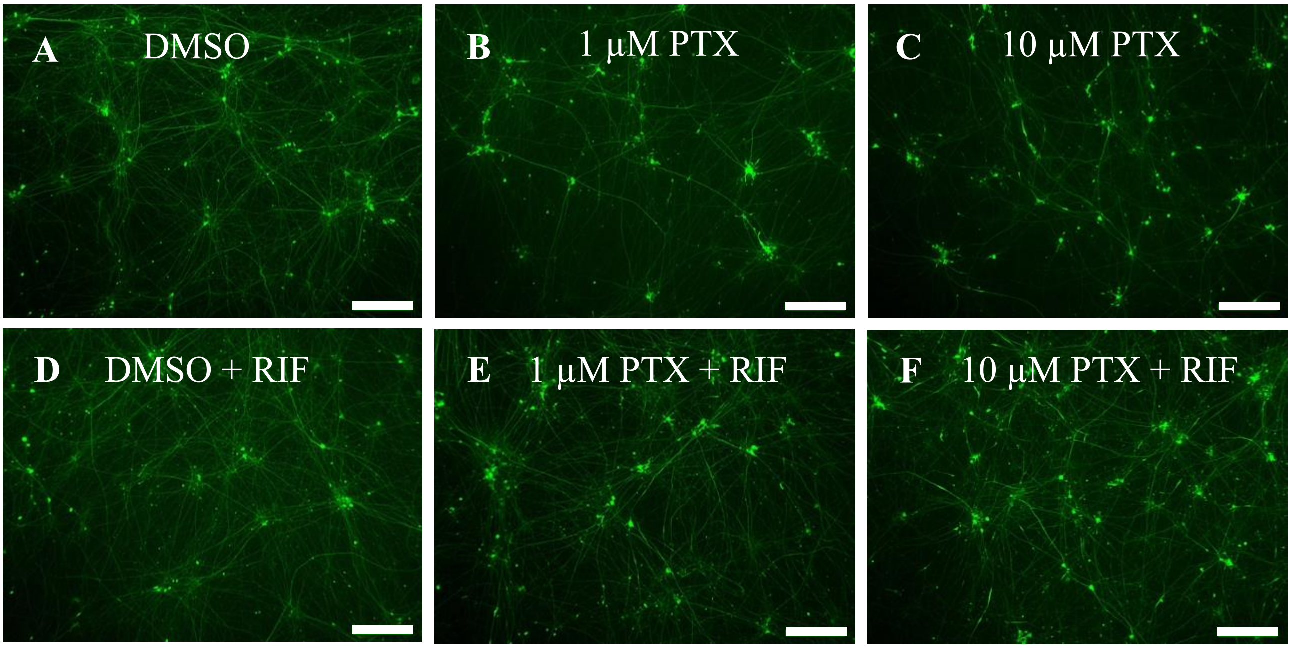


**Figure S11.** Rifampicin pre-exposure preserved the neuronal network of iPSC-SNs during paclitaxel exposure. iPSC-SNs were treated with DMSO, 1 µM and 10 µM paclitaxel with or without 48 hours of pretreatment with rifampicin. Scale bars represent 200 µm.

Abbreviations: DMSO, dimethyl sulfoxide; iPSC-SNs, induced pluripotent stem cell-derived sensory neurons; RIF, rifampicin; PTX, paclitaxel.

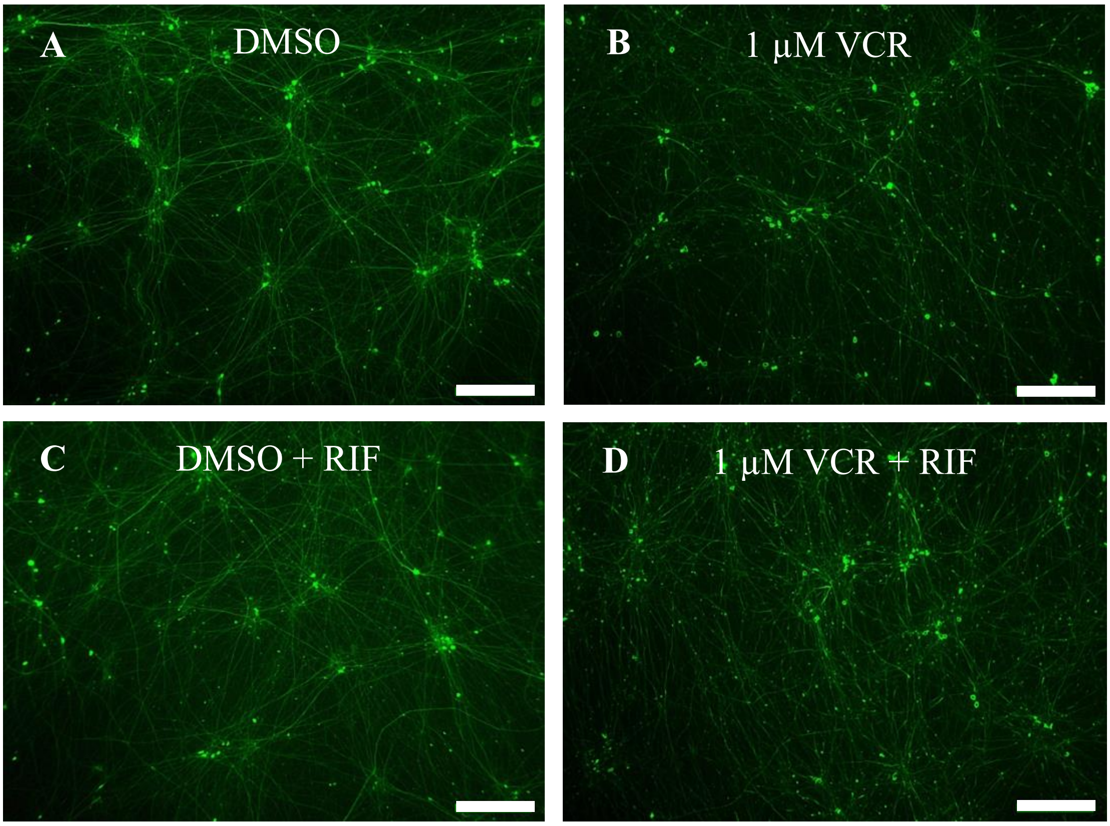


**Figure S12.** Rifampicin pre-exposure preserved the neuronal network of iPSC-SNs during vincristine exposure. iPSC-SNs were treated with DMSO and 1 μM vincristine with or without 48 hours of pretreatment with rifampicin. Scale bar represents 200 μm.

Abbreviations: DMSO, dimethyl sulfoxide; iPSC-SNs, induced pluripotent stem cell-derived sensory neurons; RIF, rifampicin; VCR, vincristine

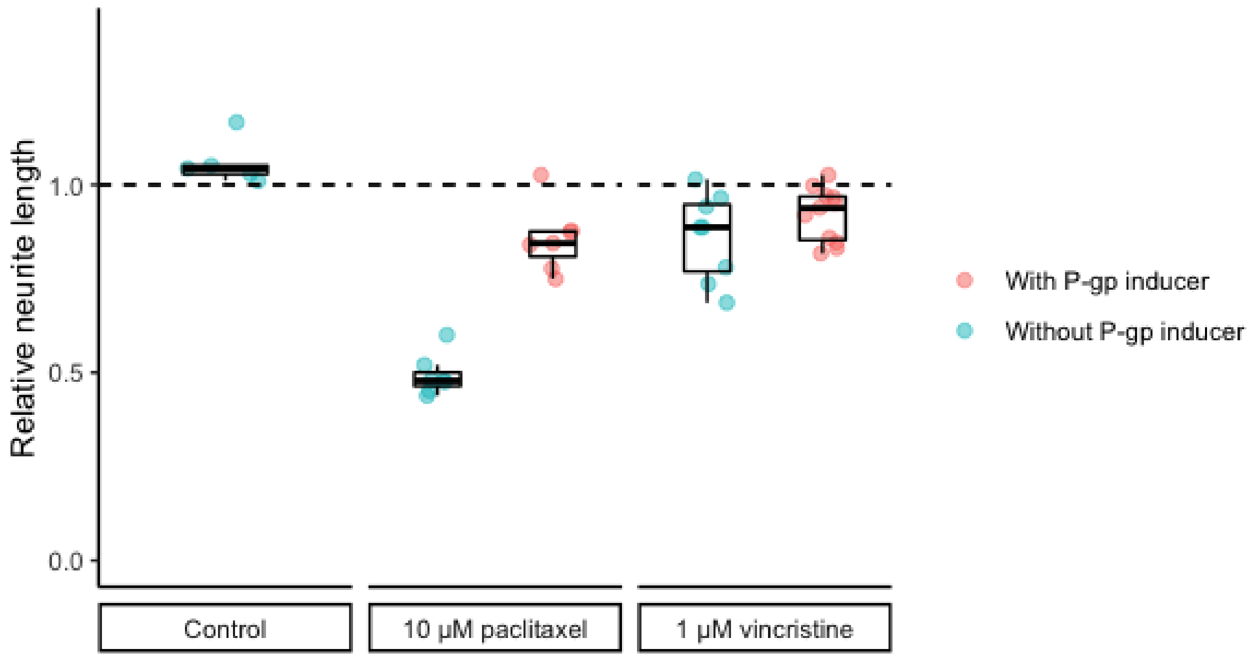


**Figure S13.** Rifampicin pre-exposure prevented both paclitaxel and vincristine neurotoxicity in iPSC-SNs. On day 38 of differentiation, iPSC-SNs were pretreated for 48 hours with rifampicin followed by 48 hours of treatment with indicated concentrations of paclitaxel or vincristine. Cells were immunolabeling with peripherin, and images were used for quantification in MIPAR. The results are presented relative to vehicle control (n=1).

Abbreviations: DMSO, dimethyl sulfoxide; iPSC-SNs, induced pluripotent stem cell-derived sensory neurons; RIF, rifampicin; VCR, vincristine.

**A**

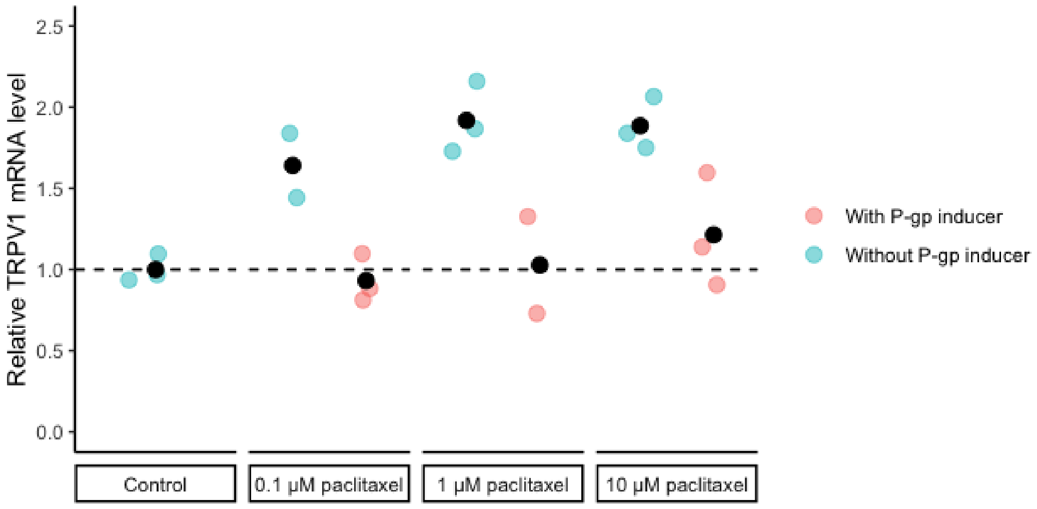


**B**

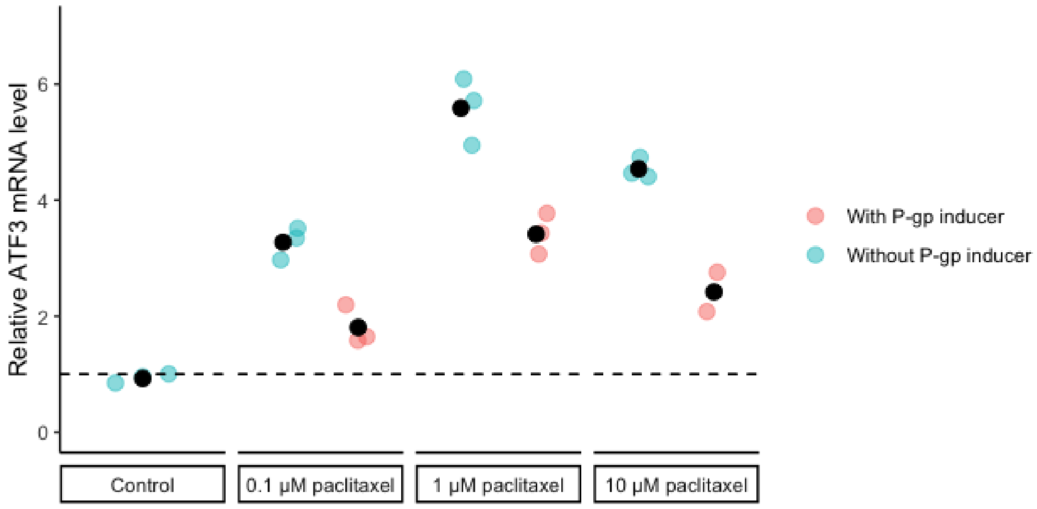


**Figure S14.** Rifampicin pre-exposure reduced *TRPV1* and *ATF3* mRNA levels in iPSC-SNs compared to paclitaxel alone. After 48 hours of induction with rifampicin iPSC-SNs were exposed to 0.01 µM, 0.1 µM, 1 µM and 10 µM PTX for 48 hours after which RNA was extracted and reverse transcribed to cDNA. The mRNA level is presented relative to the DMSO control (n=1). Abbreviations: DMSO, dimethyl sulfoxide; iPSC-SNs, induced pluripotent stem cell-derived sensory neurons; PTX, paclitaxel; RIF, rifampicin.

**Table S1.** Target-specific TaqMan probes for qPCR.

| Gene | Assay ID |
|---|---|
| GAPDH | Hs02758991_g1 |
| ATF3 | Hs00231069_m1 |
| TRPV1 | Hs00218912_m1 |
| ABCB1 | Hs00184500_m1 |
| ABCC1 | Hs01561502_m1 |

Abbreviations: ABC, ATP-binding cassette; ATF3, activating transcription factor 3; qPCR, quantitative real-time polymerase chain reaction; TRPV1, transient receptor potential vanilloid.

**Table S2**. Protein expression of ABC and SLC transporters relevant for the disposition of chemotherapy in iPSC-derived sensory neurons.

| | **Expression of ABC transporters (fmol/µg protein)** | | | | | **Expression of SLC transporters (fmol/µg protein)** | | |
|---|---|---|---|---|---|---|---|---|
| **Sample** | **P-gp (ABCB1)** | **MRP1 (ABCC1)** | **MRP2 (ABCC2)** | **MRP5 (ABCC5)** | **BCRP (ABCG2)** | **OATP1B1 (SLCO1B1)** | **OCT1 (SLC22A1)** | **OAT1 (SLC22A6)** |
| iPSC-SNs 1 | 0.02 | 0.1 | <LLOQ | <LLOQ | 0.09 | <LLOQ | <LLOQ | <LLOQ |
| iPSC-SNs 2 | 0.02 | 0.11 | <LLOQ | <LLOQ | 0.11 | <LLOQ | <LLOQ | <LLOQ |
| iPSC-SNs 3 | 0.03 | 0.14 | <LLOQ | 0.24 | 0.13 | <LLOQ | <LLOQ | <LLOQ |

Abbreviations: ABC, ATP-binding cassette, iPSC-SN, induced pluripotent stem cell-derived sensory neurons, LLOQ, lower limit of quantification; MRP, multidrug resistance-associated protein; SLC, solute carrier.
